# The RAF oncogenes of vertebrates are ohnologs that derive from the two rounds of whole genome duplications early in vertebrate evolution

**DOI:** 10.1101/767913

**Authors:** Amanda Coward Black, Mary Clay Bailey, Marisa Ruane-Foster, Juan C. Opazo, Federico G. Hoffmann

## Abstract

The Rapidly Accelerated Fibrosarcoma (RAF) kinases are part of large group of serine/threonine-specific protein kinases that play important roles in cell differentiation and organism development. Animal RAF kinases are key connectors in the signaling cascade that links the small G protein RAS and the Mitogen-activated protein kinase phosphorylation pathway. Mutations in the RAF genes have been linked to a number of cancers including melanoma, lung cancer, colorectal cancer, thyroid cancer, and ovarian cancer. Most animals possess a single RAF gene, but vertebrates have three RAF genes in their genomes, named as A-, B-, and C-RAF, the latter also known as RAF-1. The emergence of the multiple copies of vertebrate RAFs is not well resolved, on the one hand, because of sequence and functional similarities, some authors speculate that vertebrate B-Raf is most closely related to the RAF genes of the fruit fly and *Caenorhabditis elegans*, whereas a competing hypothesis is that the A-, B- and C-RAF paralogs emerged from the two rounds of whole genome duplications that occurred early in the evolution of vertebrates. We applied a comparative approach grounded in synteny and phylogenetic analyses to evaluate these two scenarios. Our results are consistent with the hypothesis that the RAF genes of vertebrates are paralogs generated by whole genome duplications. Thus, the functional similarities between vertebrate B-RAF and invertebrate RAF probably reflect the retention of ancestral characters. Interestingly, data from the literature indicate that B-RAF is the paralog that associated with cancer more strongly, and also yields the most severe phenotypes when knocked out.

## INTRODUCTION

The Rapidly Accelerated Fibrosarcoma (RAF) kinases are part of large group of serine/threonine-specific protein kinases with regulatory and signaling roles (Cseh et al. 2014). In animals, RAF kinases play important roles in cell differentiation and organism development (Desideri et al. 2015). Animal RAF kinases are key connectors in the signaling cascade that links the small G protein RAS and the Mitogen-activated protein kinase (MAPK) phosphorylation pathway (Cseh et al. 2014; Desideri et al. 2015). This pathway relays extracellular signals from receptors on the cell membrane to the cytosol to induce a variety of cellular responses, including cell proliferation, migration, differentiation, and survival (Matallanas et al. 2011; Desideri et al. 2015). Basically, RAS binds RAF to activate it, RAF then phosphorylates MEK, which then phosphorylates ERK, which can then phosphorylate a large number of downstream targets. The genes encoding for proteins in the MAPK cascade are a common source of oncogenic mutations, and the RAF genes are no exception: In fact, the first RAF gene was described as an oncogene transduced by a murine sarcoma retrovirus (Rapp et al. 1983). Later, mutations in the RAF genes have been linked to a number of cancers including melanoma, lung cancer, colorectal cancer, thyroid cancer, and ovarian cancer (Leicht et al. 2007; Desideri et al. 2015).

While most animals possess a single RAF gene, vertebrates have three RAF genes in their genomes, named as A-, B-, and C-RAF, the latter also known as RAF-1. The three vertebrate RAFs are involved in the MAPK pathway, and even though they exhibit some functional overlap, experimental evidence indicates they have independent functions and non-MAPK pathway roles as well (Wojnowski et al. 2000; Cseh et al. 2014). These three RAF isoforms can assemble into homo- or heterodimers, which expand the signaling output of the pathway in vertebrates relative to invertebrates. The emergence of the multiple copies of vertebrate RAFs is not well resolved from an evolutionary standpoint, with competing explanations about their origin. In some cases, vertebrate RAFs have been considered as putative ohnologs derived from the two rounds of whole genome duplications early in vertebrate evolution (Singh et al. 2012). On the other hand, because of sequence and functional similarities between vertebrate B-Raf and the RAF genes of the fruit fly *Drosophila melanogaster* (known as D-Raf) and the nematode *Caenorhabditis elegans* (known as lin-45), some authors suggest that B-RAF is most closely related to invertebrate RAFs than to the A- and C- paralogs of vertebrates (Matallanas et al. 2011; Desideri et al. 2015). These two hypotheses make alternative phylogenetic predictions - with vertebrate RAF paralogs expected to fall in a monophyletic clade in the first hypothesis and vertebrate B-RAF expected to group with the D-RAF gene of fruit fly and lin-45 gene of *C. elegans* in the second one - but the corresponding studies did not use phylogenetic inference to test their assertions and based their inferences on sequence similarity. Accordingly, the goals of our study are to characterize the RAF repertoire in vertebrates by integrating phylogenetic and synteny analyses to contrast competing hypotheses about the duplicative history of these genes and to incorporate an evolutionary perspective into the study of cancer-associated mutations in these genes.

## MATERIALS AND METHODS

### Sequence data

We used bioinformatic protocols to collect the repertoire of RAF-like genes in a representative set of invertebrates and vertebrates to explore copy number variation in this gene family and resolve the duplicative history of vertebrate RAFs. In the case of vertebrates, our sampling includes cyclostomes (Arctic lamprey, *Lethenteron camtschaticum*; sea lamprey, *Petromyzon marinus*; and inshore hagfish, *Eptatretus burgeri*); cartilaginous fishes (elephant shark, *Callorhinchus milii*; and whale shark, *Rhincodon typus*); and a representative sample of bony vertebrates. Bony vertebrates (Euteleostomes) were represented by one holostean fish (spotted gar, *Lepisosteus oculatus*), three teleost fish (zebrafish, *Danio rerio*; medaka, *Oryzias latipes*; Mexican cave fish, *Astyanax mexicanus*), one lobe-finned fish (West Indian Ocean coelacanth, *Latimeria chalumnae*), one amphibian (western clawed frog, *Silurana tropicalis*), four sauropsids (chicken, *Gallus gallus*; Chinese softshell turtle, *Pelodiscus sinensis*; American alligator, *Alligator mississippiensis*; and green anole lizard, *Anolis carolinensis*), and two mammals (human, *Homo sapiens*; mouse, *Mus musculus*). The search protocol combined information from the Ensembl comparative genomics assignments of orthology (Zerbino et al. 2018) with results of BLAST searches against the genomes of the American alligator, Arctic lamprey, elephant shark and whale shark.

In the case of invertebrates, we verified that the RAF genes of invertebrates are single-copy genes by screening the 70 species available in release 40 of the Ensembl Metazoa database (available at http://jul2018-metazoa.ensembl.org/index.html, last accessed on January 2019). In addition, we screened the genomes of the additional invertebrate deuterostomes- acorn worm, sea urchin, starfish and amphioxus- which are available in the NCBI sequence repository but not in Ensembl. We then selected a small number of invertebrate sequences to be included in the phylogenetic analysis corresponding to a sponge (*Amphimedon queenslandica*), a sea anemone (*Nematostella vectensis)*, a placozoan (*Trichoplax*), an arthropod (fruit fly), a nematode (*C. elegans*), two sea squirts (*Ciona intestinalis* and *Ciona savignyi*), an amphioxus (*Branchiostoma belcheri*), an acorn worm (*Saccoglossus kowalevskii*), sea urchin (*Strongylocentrotus purpuratus*), and a starfish (*Acanthaster planci*).

### Phylogenetic analyses

We inferred phylogenetic relationships among the RAF genes using the conceptual translation of the coding sequence of a representative set of the genes identified above. Sequences from the animal Kinase Suppressor of Ras (KSR) gene family, which is the animal gene family closest to RAF according to Ensembl Compara, plus a small set of plant RAF kinases were included as outgroup sequences. The full set of sequences used in the analyses are listed in supplementary table 1. We aligned amino acid sequences using Kalign (Lassmann and Sonnhammer 2006), the E-INS-i, L-INS-i, and G-INS-i strategies from MAFFT (Katoh et al. 2009; Katoh and Standley 2013), MUSCLE (Edgar 2004) and T-coffee (Notredame et al. 2000), and compared the resulting alignments using MUMSA (Lassmann and Sonnhammer 2005, 2006). Subsequently, we used the best-scoring alignment for all downstream analyses. Phylogenetic relationships were estimated using maximum likelihood (ML) and Bayesian analyses (BA). ML analyses were run using IQ-Tree ver 1.6.6 (Nguyen et al. 2015) in the implementation available from the IQ-Tree web server (Trifinopoulos et al. 2016) last accessed on September 2018, and support for the nodes was evaluated with the Shimodaira-Hasegawa approximate likelihood-ratio test (SH-aLRT), the aBayes test (Anisimova et al. 2011) and 1,000 pseudoreplicates of the ultrafast bootstrap procedure (Hoang et al. 2018). The best-fitting model of substitution was selected using the ModelFinder subroutine from IQ-Tree (Kalyaanamoorthy et al. 2017). Bayesian Analyses were performed via the CIPRES Science Gateway (Miller et al. 2015) in MrBayes version 3.2 (Ronquist et al. 2012), under a mixed model of substitution, running four simultaneous chains for 2 × 10^7^ generations, sampling trees every 1000 generations, and using default priors. We assessed convergence by measuring the standard deviation of the split frequency among parallel chains. Chains were considered to have converged once the average split frequency was lower than 0.01. We discarded trees collected before the chains reached convergence, and we summarized results with a majority-rule consensus of trees collected after convergence was reached.

### Synteny comparisons

Whole-genome duplications are expected to affect all genes in a given genome. So, synteny analyses can help identify sets of co-duplicated genes that map to similar regions of the genome. To do so, we examined genes found upstream and downstream of each member of the RAF gene family on representative species. Initial ortholog predictions were derived from the EnsemblCompara database (Zerbino et al. 2018) and visualized using the Genomicus platform v92.01, last accessed in August 2018 (Nguyen et al. 2018). In the case of the elephant shark, synteny was visualized on NCBI and inferred by comparing flanking genes to other vertebrates using BLAST tools.

### Organismal phylogeny and divergence dates

In all cases, we assumed that relationships among the different animal lineages follow the arrangement reported by (Dunn et al. 2014), and estimates of divergence times among the lineages were obtained from the TimeTree server (Kumar et al. 2017).

## RESULTS AND DISCUSSION

### RAF repertoires

We found limited variation in the RAF repertoires of animals. In the case of invertebrates, our bioinformatic surveys confirm that most invertebrates include a single member of this gene family in their genome. Fifty five species in a total of 70 have a 1-to-1 ortholog of the *Drosophila melanogaster* D-Raf gene, 7 have 1-to-many orthologs, and 8 have no annotated ortholog. A more critical evaluation of the sequences of the 7 species that have putative duplicates indicates that all of these duplicates correspond to either very recent duplications or to assembly artifacts. Pairwise comparisons among the putative RAF homologs within a species are either identical; as in the stalk-eyed fly *Teleopsis dalmanni*, the nematodes *Caenorhabditis brenneri* and *Caenorhabditis remanei*, the rotifer *Adineta vaga*, the cnidarian *Nematostella vectensis*, and the brachiopod *Lingula anatina*; or correspond to two fragments of a single gene, as in the case of the African social velvet spider *Stegodyphus mimosarum*. Bioinformatic searches in other invertebrate genomes in other repositories were consistent with these results and identified a single putative RAF ortholog in most cases, either spanning the gene completely or partially, as in the case of the amphioxus *Branchiostoma floridae*. Because we found RAF orthologs in a poriferan (the sponge *Amphimedon queenslandica*), a ctenophore (the warty comb jelly *Mnemiopsis leidyi*), the placozoan *Trichoplax adherens*, a cnidarian (the sea anemone *Nematostella vectencis*), protostomes, and deuterostomes, we can trace the presence of the animal RAF gene back to the common ancestor of animals, though we could not find candidates in choanoflagellates.

In the case of vertebrates, most species include three RAF-like genes in their genomes, A-, B- and C-Raf, with the exception of the two lampreys, which include one or two paralogs, and Archelosaurs (the group that includes birds, crocodilians, and testudines), where we could find copies of B- and C-RAF, but not of A-RAF. Two of the teleost fish surveyed, zebrafish and Mexican cave fish, possess duplicate copies of C-RAF, and there are 2 separate fragments of A-RAF in the JGI4.2/XenTro3 release of the Western clawed frog (*Silurana tropicalis*) genome, which are located on separate scaffolds and annotated as separate genes. This is probably an assembly artifact, as these two fragments map to non-overlapping portions of the human A-RAF gene. We included one of the A-RAF homologs of the closely related African clawed frog (*Xenopus laevis*) genome in the analyses as an additional representative of amphibians. The current assembly of the elephant shark genome lacks an A-Raf gene, but a partial match is found in the whale shark genome, indicating the gene was present in the ancestor of cartilaginous fish.

From a sequence standpoint, pairwise distance comparisons within the A-, B-, and C-paralogs of jawed vertebrates reveal that the B-RAF paralog is the most conserved, the C-RAF paralog is strongly conserved as well, and the A-RAF paralog is the most variable (Table 1)

**Table 1.**
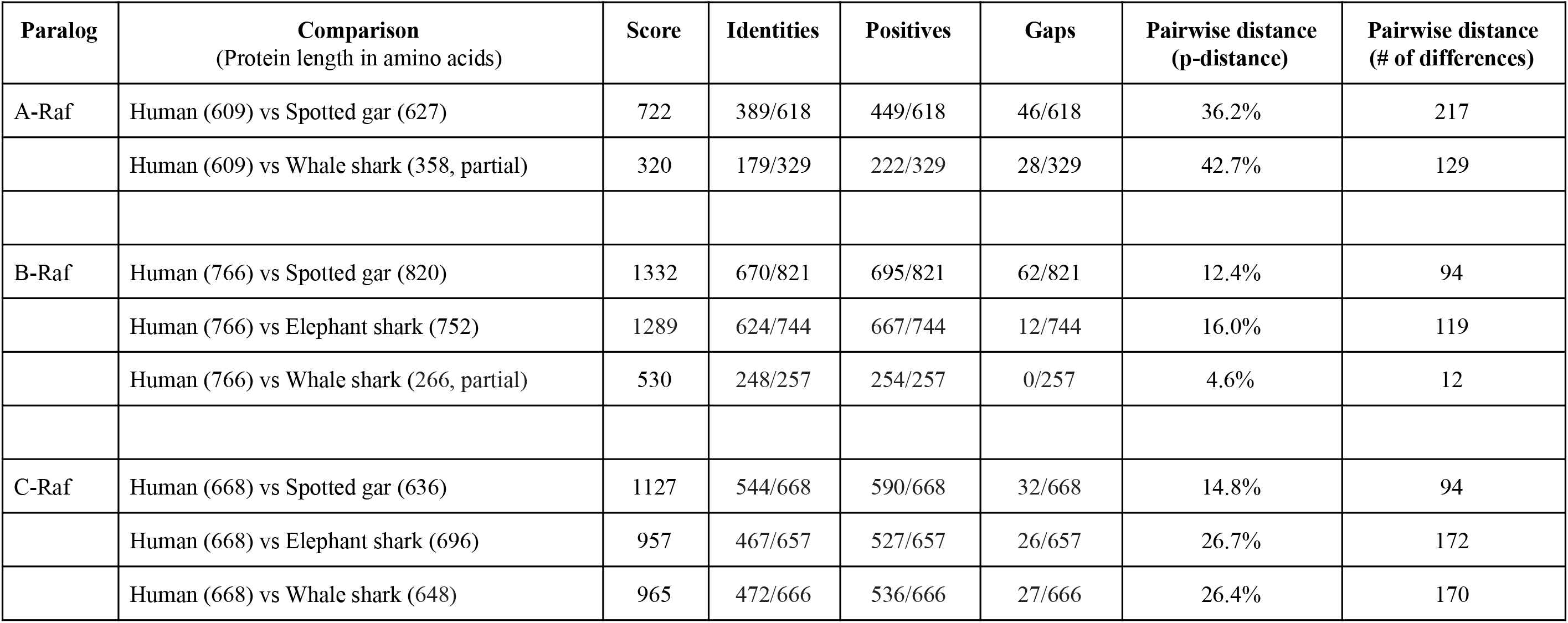
Pairwise comparisons between the A-,B-, and C-Raf paralogs of human and spotted gar, and those of human and elephant or whale shark.

### Evolutionary history of the RAF oncogenes

Our phylogenetic analyses include a representative subset of animal RAFs, plus a small sample of plant RAF-like and animal KSR sequences as outgroup sequences (see Supplementary Table 1 for a full list of sequences used in phylogeny reconstructions). The resulting trees show that animal RAFs form a monophyletic group that is distantly related to plant RAFs or animal KSR sequences (Fig. 1). Relationships between plant and animal RAFs are poorly understood and their functional roles seem to be different: plant RAFs are mostly involved with signaling networks involved in responses to stress, whereas animal RAFs are involved with cell cycle progression, cell differentiation, migration and organismal development (Popescu and Popescu 2011; Lehti-Shiu and Shiu 2012). Sequence similarities between animal and plant RAFs are restricted to the kinase domain. In agreement with these observations, support for the node uniting animal and plant RAFs is very low. Animal RAFs are separated into two distinct groups, one including vertebrate RAFs and the other one including invertebrate RAFs (Fig. 1). This arrangement is not entirely consistent with the expected organismal relationships, as it does not group all deuterostomes in a monophyletic group, however, support for the relevant nodes is weak. We suspect that deep divergences among some invertebrates in our study, potential inconsistencies in the annotations of the gene models, and incomplete coverage for some genes in our study could account for this observation.

**Figure 1.**
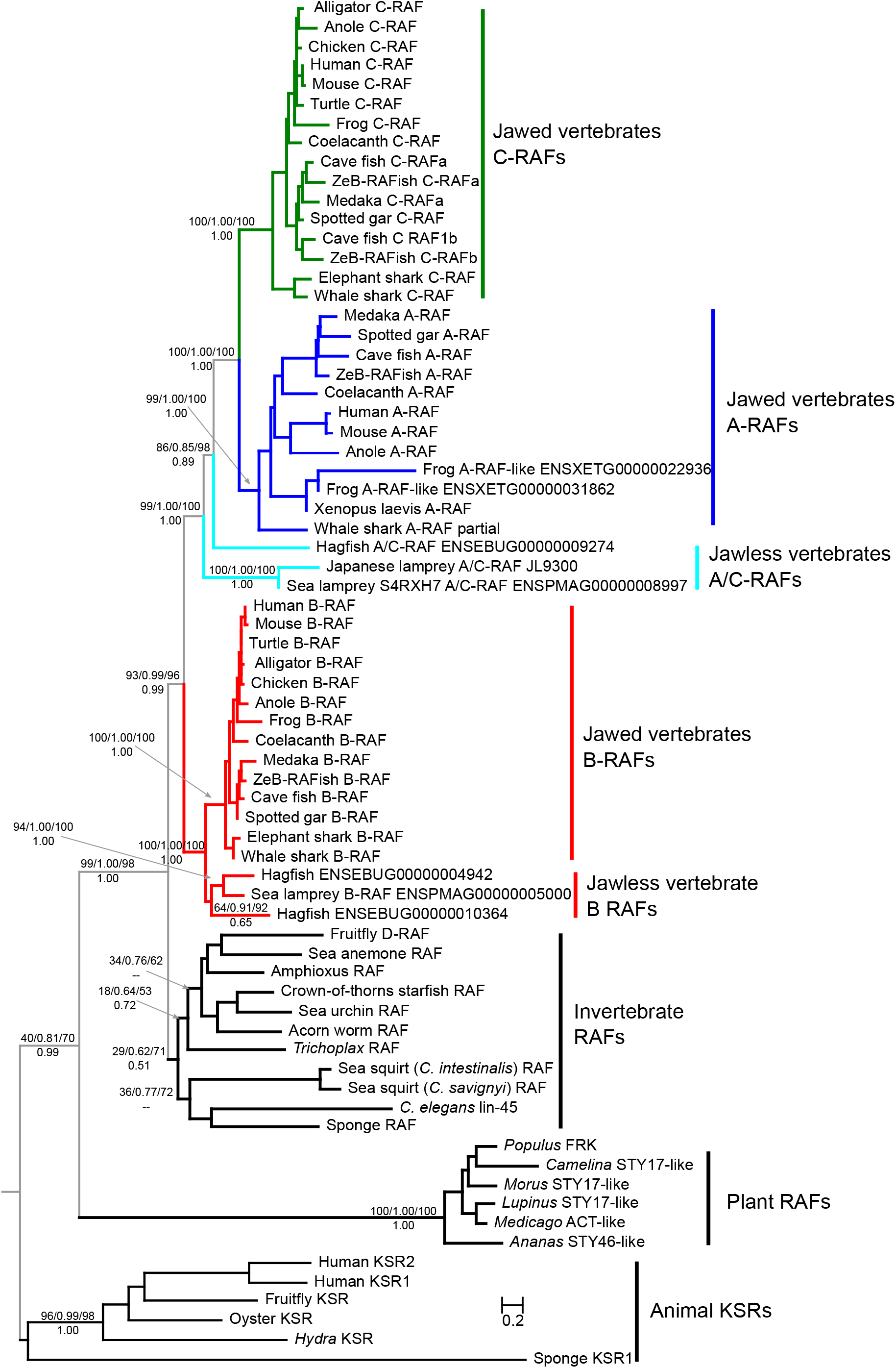
Maximum-likelihood phylogram depicting evolutionary relationships among animal RAFs, with plant RAFs and animal KSRs as outgroup sequences. Support for the relevant nodes is shown next to the relevant nodes. Numbers above the nodes correspond to support from the Shimodaira-Hasegawa approximate likelihood-ratio test/ aBayes/ maximum likelihood ultrafast bootstrap support, and number below the nodes correspond to Bayesian posterior probability support values. The coloring of vertebrate RAFs reflects their orthology: B-RAFs in red, jawed vertebrates C-RAFs in green, and A-RAFs in blue, and the A/C RAFs of jawless vertebrates in light blue. Note that the A/C RAFs of lampreys and hagfish are not monophyletic and might represent paralogous genes.

In both our ML and BA phylogenies, vertebrate RAFs are placed in a monophyletic group with strong support (93/0.99/96/1.0), and these analyses resolved orthology and duplicative history for jawed vertebrate RAFs well (Fig. 1). Each of the A-, B- and C-RAF genes of jawed vertebrates are recovered in a strongly supported monophyletic group, with the A- and C-RAF paralogs as sister groups, and the B-RAF paralog as the deepest split within vertebrate RAFs (Fig. 1). Relationships among the genes within each of these clades did not deviate significantly from the expected organismal tree, placing cartilaginous fish sequences as sister to all other jawed vertebrates, and grouping fish and tetrapod paralogs in monophyletic groups within each RAF paralog clade (Fig. 1). On the other hand, our analyses could not fully resolve orthology for the RAF sequences from cyclostomes. This is not surprising, as resolving orthology between gnathostome and cyclostome genes using phylogenies has had limited (Qiu et al. 2011; Kuraku 2013; Schwarze et al. 2014; Opazo et al. 2015; Campanini et al. 2015). In the case of B-RAF, our analyses identified one putative copy of this gene in the sea lamprey and two in the hagfish. The tree places the sea lamprey gene as sister to one of the hagfish B-RAF paralogs, a phylogenetic arrangement that would suggest that the last common ancestor of lamprey and hagfish had duplicate copies of B-RAF in its genome. Lampreys and hagfish include an additional RAF gene with affinities for the A/C-RAF lineage of jawed vertebrates, but they have no clear orthologs. The hagfish A/C-RAF gene is placed sister to the clade that includes the RAF A- and C- paralogs of jawed vertebrates, and the A/C-RAF genes of lampreys fall in a clade sister to the group that unites the hagfish A/C-RAF gene and the RAF-A and -C paralogs of jawed vertebrates (Fig. 1).

### RAF synteny

Synteny comparisons are consistent with the results of the pairwise sequence comparisons and phylogenetic analyses. We observed higher synteny conservation in the B- and C-RAF genes of jawed vertebrates relative to the A-RAF paralog (Supplementary Fig. 1). In the case of A-Raf, there are very few genes that are syntenic between gar and human, which diverged approximately 430 million years ago. By contrast, in the case of C-RAF, the 6 genes found downstream are conserved in comparisons all the way from human to elephant shark, which diverged approximately 470 million years ago, and 4 genes upstream are conserved from anole to elephant shark. In addition, one of the two zebrafish C-RAF duplicates also shows some conservation with the genes upstream from human C-RAF. The B-RAF gene shows an intermediate pattern, with strong conservation in amniotes, as represented by comparisons between anole and human, which diverged approximately 318 million years ago, but more limited in more distant comparisons. Still, there are copies of MRPS33 and TMEM178B immediately upstream of B-RAF in most jawed vertebrates assessed, plus a copy of WEE2 that is located downstream in the elephant shark and upstream in amniotes (Fig. 2). Unfortunately, synteny provides limited information to help resolve orthology between cyclostome and gnathostome RAFs, as the RAF paralogs of the sea lamprey and hagfish have very limited information on flanking genes: the A/C-RAF of hagfish is flanked by orthologs of RAB43 and ISY1 which are flanking C-RAF in jawed vertebrates.

**Figure 2.**
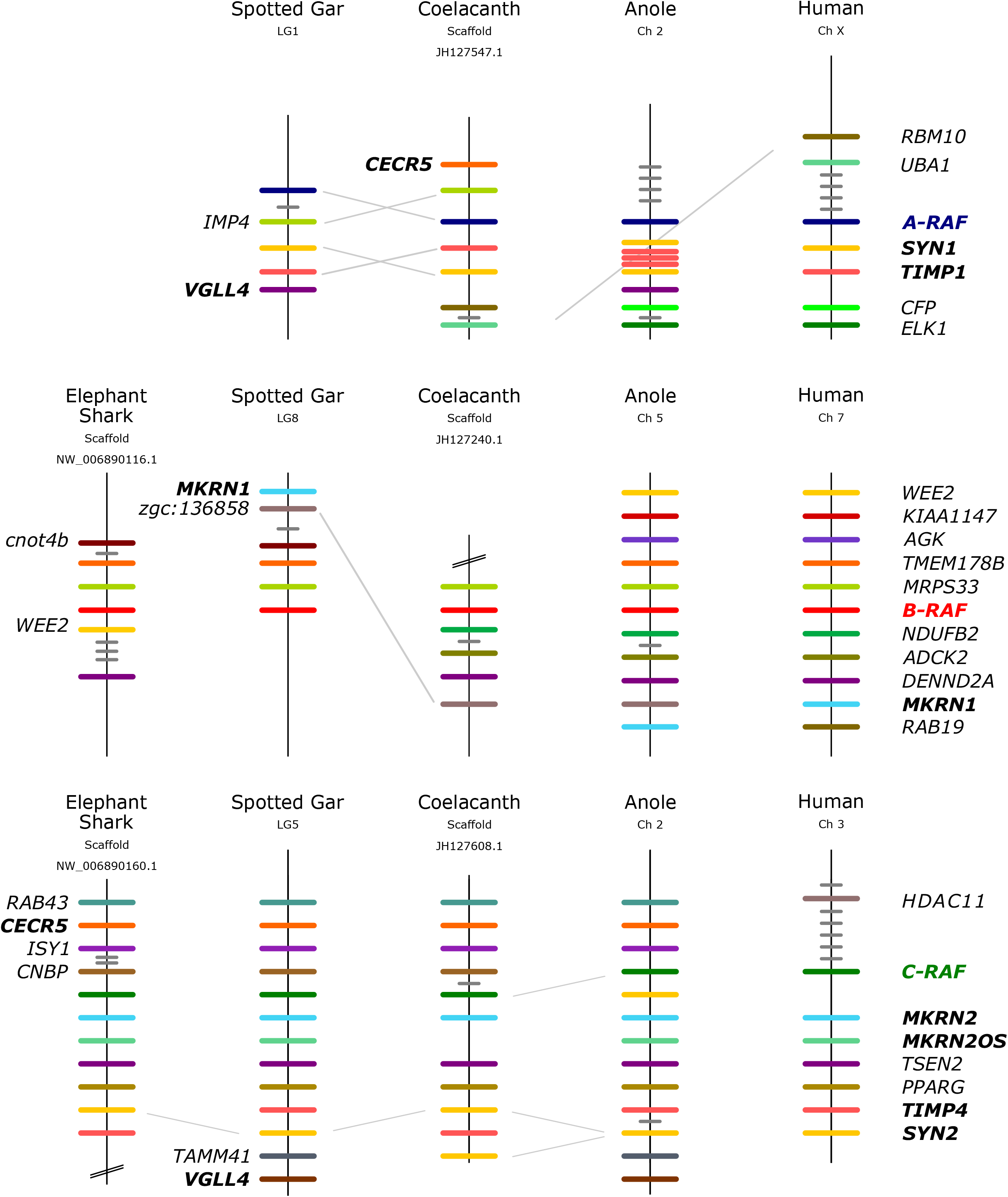
Patterns of conserved synteny in genomic regions that harbor paralogous RAF genes in representative vertebrate taxa. Genes are color coded following orthology. Genes that co-duplicated along with the RAF paralogs are in bold. Diagonal lines connecting genes indicate a change of position.

### Origin of vertebrate RAFs

The two competing hypotheses about the emergence of animal RAFs make alternative phylogenetic predictions. If the B-RAF of vertebrates was more closely related to the single copy RAFs of *C. elegans* and fruit fly, these genes would fall in a monophyletic group to the exclusion of the other RAF paralogs of vertebrates, and this is clearly not that case (Fig. 1). Alternatively, if vertebrate RAFs were ohnologs that expanded as a result of the two rounds of whole genome duplications in early vertebrate evolution, they would be monophyletic relative to invertebrate RAFs, in agreement with our phylogenies (Fig. 1). In addition, synteny comparisons reveal that the genomic context of the A-, B-, and C-RAF paralogs is similar in most species of jawed vertebrates analyzed, which would indicate that their location in the genome was established early in vertebrate evolution and has remained relatively conserved (Fig. 2). The human A-, B-, and C-RAF paralogs are located on chromosomal segments X.4, 7.6 and 3.1 respectively, which all map to ancestral Chordate Linkage Group 13 (Putnam et al. 2008). Thus, our results are consistent with the hypothesis that the RAF genes of jawed vertebrates are ohnologs derived from the two rounds of WGD early in vertebrate evolution, and the presence of co-duplicated genes MKRN, CECR5, VGLL4, synapsin, and TIMP gene families linking the different RAF paralogons is also in line with this inference. Interestingly, the A-, B- and C-RAF genes appear to have reverted to their single-copy state in most teleosts after the whole genome duplication occurred in that lineage, with the exception of the C-RAF gene in zebrafish and Mexican cave fish.

The three RAFs of jawed vertebrates work by phosphorylating MEK, with some MEK-independent pathways involving apoptosis (Desideri et al. 2015). All three show some functional redundancy, as evidenced by RAF knockouts and a comparison of single and double B- and C-RAF mutants (Wojnowski et al. 2000; Galabova-Kovacs et al. 2006). In the case of B-RAF, transgenic mice knockouts died in utero by embryonic day 10-12.5 (Wojnowski et al. 1997). In the case of C-RAF, homozygous knockouts died by embryonic day 12.5, while heterozygous died shortly after birth (Wojnowski et al. 1998). In the case of A-RAF, knockout mice of the C57Bl/6J strain die 7-21 days after birth while knockouts in mice outbred with the 129/OLA strain survive to adulthood with some neurological defects (Pritchard et al. 1996). Thus, the loss of B- or C-RAF appears to have more severe consequences relative to A-RAF, an inference that is in line with the fact that archelosaurs live without a copy of this gene, but that B- and C-RAF are present in all vertebrates. This is also in line with known essential roles of both B- and C-RAF, while most characterized roles of A-RAF overlap with the other two (Cseh et al. 2014; Desideri et al. 2015).

### Gene copy retention and cancer

Utilizing the OncoKB database (Chakravarty et al. 2017), we identified 138 annotated point mutations and exon deletions across all three RAFs that are cancer-associated mutations or sites examined within the context of carcinogenic properties (Table 2). We found that most disruptive mutations map to the kinase domains of the B-RAF paralog, where 76% of confirmed oncogenic mutations and 61% of likely oncogenic mutations are found. Additionally, of the 138 alterations, all but 20 have an effect that is gain-of-function or likely gain-of-function and only 4 have an effect that is loss-of-function or likely loss-of-function (Supplementary Table 2). Singh et al. (2012) classified putative ohnologs derived from the 2R of WGD of vertebrates as either ‘essential’, defined as genes where loss-of-function mutations have very severe phenotypic consequences, or ‘dangerous’, defined as genes where gain of function mutations lead to constitutive activity and are in turn linked to cancer (Singh et al. 2012). They then proposed a model where the retention of ohnologs was mostly driven by ‘their “dangerousness.”’ Our results align well with this prediction. Further, if we take a more expansive definition of essentiality, where in addition to loss-of-function we also consider the phenotypic consequences of the loss of a given gene, both essentiality and dangerousness would align. B-RAF is the one that exhibits more severe phenotypes when lost, closely followed by C-RAF. Similarly, most cancer mutations in the gene family map to the B-RAF paralog followed by C-RAF, which are the two most conserved paralogs in terms of sequence, synteny, and phyletic distribution.

**Table 2.**
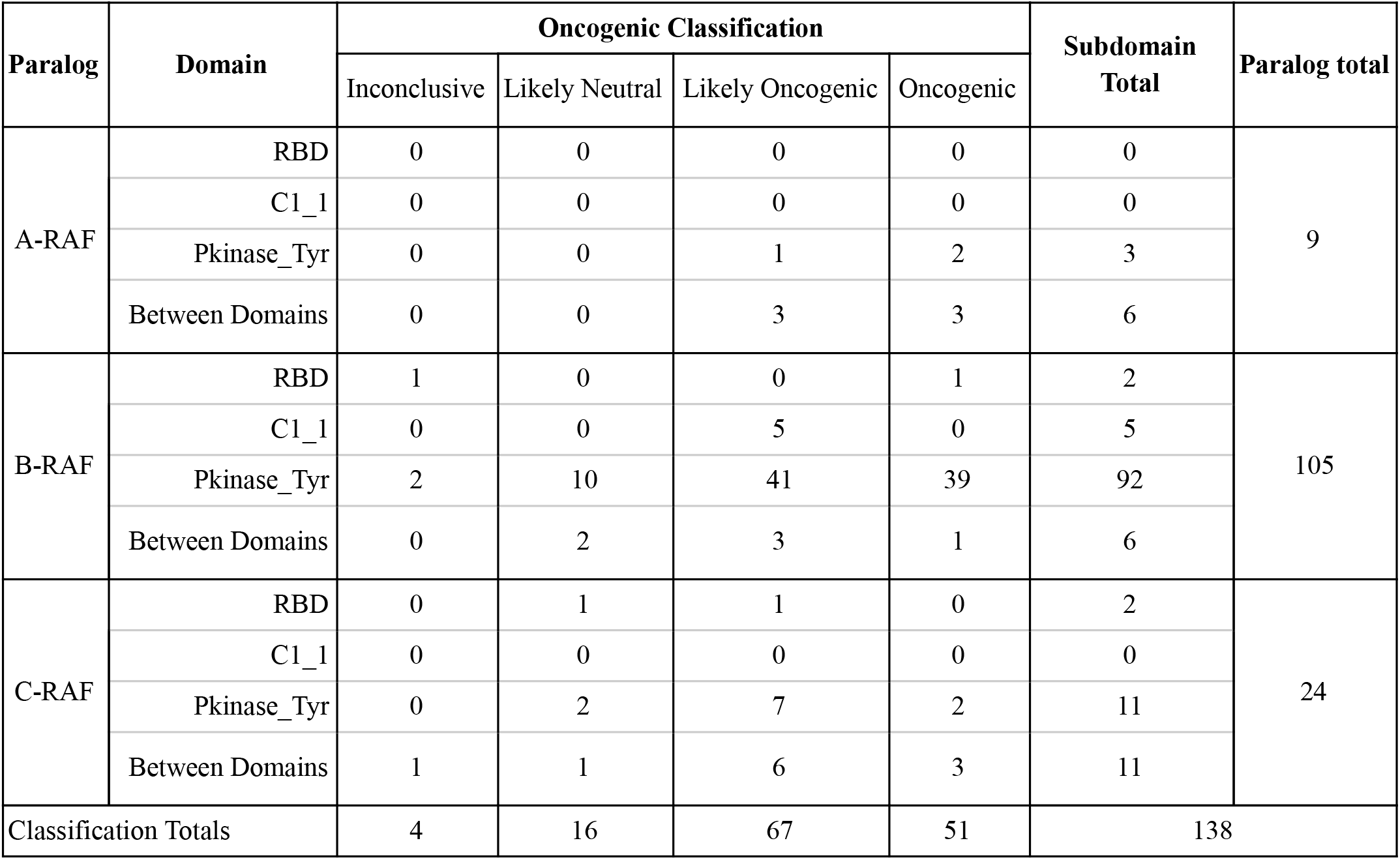
Summary of OncoKB annotated mutations in RAF paralogs.

| Paralog | Domain | Oncogenic Classification |  |  |  | Subdomain Total | Paralog total |
| --- | --- | --- | --- | --- | --- | --- | --- |
|  |  | Inconclusive | Likely Neutral | Likely Oncogenic | Oncogenic |  |  |
| A-RAF | RBD | 0 | 0 | 0 | 0 | 0 | 9 |
|  | C1_1 | 0 | 0 | 0 | 0 | 0 |  |
|  | Pkinase_Tyr | 0 | 0 | 1 | 2 | 3 |  |
|  | Between Domains | 0 | 0 | 3 | 3 | 6 |  |
| B-RAF | RBD | 1 | 0 | 0 | 1 | 2 | 105 |
|  | C1_1 | 0 | 0 | 5 | 0 | 5 |  |
|  | Pkinase_Tyr | 2 | 10 | 41 | 39 | 92 |  |
|  | Between Domains | 0 | 2 | 3 | 1 | 6 |  |
| C-RAF | RBD | 0 | 1 | 1 | 0 | 2 | 24 |
|  | C1_1 | 0 | 0 | 0 | 0 | 0 |  |
|  | Pkinase_Tyr | 0 | 2 | 7 | 2 | 11 |  |
|  | Between Domains | 1 | 1 | 6 | 3 | 11 |  |
| Classification Totals |  | 4 | 16 | 67 | 51 | 138 |  |

Integrating results of our phylogenetic, synteny, and sequence conservation analyses with literature searches suggests the RAF paralogs of vertebrates derive from the 2R of WGD early in the evolution of vertebrates. Our results imply that the functional similarities between the single copy RAF gene of invertebrates and the B-RAF paralog of jawed vertebrates derive from the retention of shared ancestral states and do not reflect evolutionary origin. This is probably also true for the A-, B- and C-RAF paralogs of jawed vertebrates, where functional similarities between B- and C- are probably ancestral, and functional similarities between A- and C-RAF reflect derived functional differentiation (Matallanas et al. 2011; Desideri et al. 2015). Integrating functional data into our phylogenetic analyses would suggest that all three vertebrate RAF paralogs have maintained some of their ancestral roles, with functional divergence among them: B-RAF is the most potent activator of ERK whereas A-RAF is the least potent one (Desideri et al. 2015). However, and in contrast to the single-copy invertebrate counterparts, the different RAF proteins of vertebrates can assemble into heterodimers, expanding the potential outputs of the RAF signaling cascade, and each of them plays a slightly different role in the pathway, which suggest that their functional divergence involves a combination of subfunctionalization and neofunctionalization processes.

## ACKNOWLEDGEMENTS

We thank Sorina C. Popescu for help with identifying relevant plant RAF kinases. This work was partially funded by the National Science Foundation through grants EPS-0903787, DBI-1659630, DEB-1354147 and EPSCoR RII Track-2 FEC 1736026 to FGH, Millennium Nucleus of Ion Channels Associated Diseases (MiNICAD), Iniciativa Científica Milenio, Ministry of Economy, Development and Tourism from Chile to JCO and Fondo Nacional de Desarrollo Científico y Tecnológico from Chile (FONDECYT 1160627) to JCO.

---

Supplementary Figure 1. Expanded synteny.

Supplementary Table 1. Sequence IDs.

Supplementary Table 2. Cancer mutations.

