## Supplemental Figure 1 and Supplemental Tables 1-2 for "The RAF oncogenes of vertebrates are ohnologs that derive from the two rounds of whole genome duplications early in vertebrate evolution"

Supplemental Table 1.

| Species | Scientific Name | Gene | Lineage | Class | Accession Number | Genome Release | Source |
|---|---|---|---|---|---|---|---|
| African clawed frog | *Xenopus laevis* | A-RAF-like_ partial | Amphibia | Amphibia | NP_001083376 | xenLae2 | NCBI |
| Western clawed frog | *Xenopus tropicalis* | A-RAF | Amphibia | Amphibia | ENSXETG00000022936 | JGI 4.2 | Ensembl |
| | | A-RAF-like | | | ENSXETG00000031862 | | |
| | | B-RAF | | | ENSXETG00000004845 | | |
| | | C-RAF | | | ENSXETG00000020163 | | |
| Fruitfly | *Drosophila melanogaster* | RAF | Arthropod | Insecta | FBpp0310458 | BDGP6.22 | Ensembl |
| | | KSR | | | AAC46970 | Release 6 plus ISO1 MT | NCBI |
| Pacific oyster | *Crassostrea gigas* | KSR1 | Bivalve | Bivalvia | XP_011432744 | oyster_v9 | NCBI |
| | | KSR2 | | | XP_011432742 | | |
| Elephant shark | *Callorhinchus milii* | B-RAF | Cartilaginous fish | Chondrichthyes | 103181856 | Callorhinchus_milii-6.1.3 | NCBI |
| | | C-RAF | | | 103185022 | | |
| Whale shark | *Rhincodon typus* | A-RAF-like_ partial | Cartilaginous fish | Chondrichthyes | XP_020380753 | ASM164234v2 | NCBI |
| | | B-RAF-like_ partial | | | XP_02376592 | | |
| | | C-RAF | | | XP_020366455 | | |
| Amphioxus | *Branchiostoma belcheri* | RAF | Chordate | Leptocardii | XP_019633461 | Haploidv18h27 | NCBI |
| Solitary sea squirt | *Ciona savignyi* | H2Y816 | Chordate | Ascidiacea | ENSCSAVG00000000836 | CSAV 2.0 | Ensembl |

| | | | | | | | |
|---|---|---|---|---|---|---|---|
| Vase tunicate | *Cionia intestinalis* | C-RAF | Chordate | Ascidiacea | ENSCING00000003721 | KH | Ensembl |
| Freshwater polyp | *Hydra vulgaris* | KSR1 | Cnidaria | Hydrozoa | XP_012560417 | Hydra_RP_1.0 | NCBI |
| | | KSR2 | | | XP_012563722 | | |
| Sea anemone | *Nematostella vectensis* | NEMVEDRAFT_v1g60102 | Cnidaria | Anthozoa | XP_001639152 | NEMVEDRAFT | NCBI |
| Starfish | *Acanthaster planci* | RAF | Echinoderm | Asteroidea | XP_022104905 | OKI-Apl_1.0 | NCBI |
| Sea urchin | *Strongylocentrotus purpuratus* | RAF | Echinoderm | Echinoidea | SPU_027524 | Spur_3.1 | Ensembl |
| Acorn worm | *Saccoglossus kowalevskii* | RAF | Hemichordata | Enteropneusta | XP_006821654 | Skow_1.1 | NCBI |
| Spotted gar | *Lepisosteus oculatus* | A-RAF | Holost fish | Actinopterygii | ENSLOCG00000014384 | LepOcu1 | Ensembl |
| | | B-RAF | | | ENSLOCG00000016118 | | |
| | | C-RAF | | | ENSLOCG00000013974 | | |
| Hagfish | *Eptatretus burgeri* | B-RAF | Jawless fish | Myxini | ENSEBUG00000004942 | Eburgeri_3.2 | Ensembl |
| | | A-RAF | | | ENSEBUG00000009274 | | |
| | | B-RAF | | | ENSEBUG00000010364 | | |
| Japanese lamprey | *Lethenteron japonicum* | JL9300 | Jawless fish | Agnatha | Not available | | JL Genome Project |
| Sea lamprey | *Petromyzon marinus* | S4RXH7 | Jawless fish | Agnatha | ENSPMAG00000008997 | Pmarinus_7.0 | Ensembl |
| | | B-RAF | | | ENSPMAG00000005000 | | |
| Coelacanth | *Latimeria chalumnae* | A-RAF | Lobe-finned fish | Sarcopterygii | ENSLACG00000009313 | LatCha1 | Ensembl |
| | | B-RAF | | | ENSLACG00000011252 | | |
| | | C-RAF | | | ENSLACG00000007082 | | |

| | | | | | | | |
|---|---|---|---|---|---|---|---|
| Human | *Homo sapiens* | A-RAF | Mammal | Mammal | ENSG00000078061 | GRCh38 | Ensembl |
| | | B-RAF | | | ENSG00000157764 | | |
| | | C-RAF | | | ENSG00000132155 | | |
| | | KSR1 | | | NP_055053 | | NCBI |
| | | KSR2 | | | NP_775869 | | |
| Mouse | *Mus musculus* | A-RAF | Mammal | Mammal | ENSMUSG00000001127 | GRChm38 | Ensembl |
| | | B-RAF | | | ENSMUSG00000002413 | | |
| | | C-RAF | | | ENSMUSG00000000441 | | |
| Nematode | *Caenorhabditis elegans* | lin-45 | Nematode | Chromadorea | Y73B6A | WBcel235 | Ensembl |
| Trichoplax | *Trichoplax adhaerens* | RAF_partial | Placozoa | Placozoa | TRIADDRAFT_24886 | v1.0 (GCF_000150275.1) | NCBI |
| Barrelclover | *Medicago truncatula* | ACT-like | Plant | Eudicot | KEH27604 | MedtrA17_4.0 | NCBI |
| Desert poplar | *Populus euphratica* | FRK | Plant | Eudicot | XP_011022235 | PopEup_1.0 | NCBI |
| False flax | *Camelina sativa* | STY17-like | Plant | Eudicotidae | XP_010432231 | Cs | NCBI |
| Mulberry tree | *Morus notabilis* | STY17-like | Plant | Eudicot | XP_024032796 | ASM41409v2 | NCBI |
| Narrow-leaved blue lupine | *Lupinus angustifolius* | STY17-like | Plant | Eudicots | XP_019421297 | LupAngTanjil_v1.0 | NCBI |
| Pineapple | *Ananas comosus* | STY46-like | Plant | Monocotyledon s | XP_020112392 | ASM154086v1 | NCBI |
| Sponge | *Amphimedon queenslandica* | RAF | Porifera | Demospongiae | XP_019850658 | v1.0 (GCA_000090795.1) | NCBI |
| | | KSR1 | | | XP_019852299 | | |
| Softshell turtle | *Pelodiscus sinensis* | B-RAF | Sauropsid | Reptile | ENSPSIG00000011427 | PelSin_1.0 | Ensembl |
| | | C-RAF | | | ENSPSIG00000009677 | | |

| | | | | | | | |
|---|---|---|---|---|---|---|---|
| American Alligator | *Alligator mississippiensis* | B-RAF | Sauropsida | Reptilia | 106737102 | ASM28112v4 | NCBI |
| | | C-RAF | | | 102576986 | | |
| Anole | *Anolis carolinensis* | A-RAF | Sauropsida | Reptilia | ENSACAG00000011249 | AnoCar2.0 | Ensembl |
| | | B-RAF | | | ENSACAG00000011178 | | |
| | | C-RAF | | | ENSACAG00000013194 | | |
| Chicken | *Gallus gallus* | B-RAF | Sauropsida | Aves | ENSGALG00000012865 | Gallus_gallus-5.0 | Ensembl |
| | | C-RAF | | | ENSGALG00000004998 | | |
| Medaka | *Oryzias latipes* | A-RAF | Teleost fish | Actinopterygii | ENSORLG00000016733 | HdrR | Ensembl |
| | | B-RAF | | | ENSORLG00000009843 | | |
| | | C-RAFa | | | ENSORLG00000011471 | | |
| Mexican tetra/blind cave fish | *Astyanax mexicanus* | A-RAF | Teleost fish | Actinopterygii | ENSAMXG00000004222 | Astyanax_mexicanus -2.0 | Ensembl |
| | | B-RAF | | | ENSAMXG00000011513 | | |
| | | C-RAFa | | | ENSAMXG00000001055 | | |
| | | C-RAFb | | | ENSAMXG00000005440 | | |
| Zebra fish | *Danio rerio* | A-RAF | Teleost fish | Actinopterygii | ENSDARG00000054533 | GRCz10 | Ensembl |
| | | B-RAF | | | ENSDARG00000017661 | | |
| | | C-RAFa | | | ENSDARG00000096415 | | |
| | | C-RAFb | | | ENSDARG00000059406 | | |

Supplemental Table 2.

| Mutation | RAF Paralog | Oncogenic? | Location of Mutation | Mutation Effect | Citation |
|---|---|---|---|---|---|
| S214A | A-RAF | Likely | Between C1_1 and Tyrosine Protein Kinase | Likely Gain-of-functi on | Imielinski, M., Greulich, H., Kaplan, B., Araujo, L., Amann, J., Horn, L., … Carbone, D. P. (2014). Oncogenic and sorafenib-sensitive ARAF mutations in lung adenocarcinoma.Journal of Clinical Investigation,124(4), 1582–1586. https://doi.org/10.1172/JCI72763 |
| S214C | A-RAF | Yes | Between C1_1 and Tyrosine Protein Kinase | Gain-of-functi on | Imielinski, M., Greulich, H., Kaplan, B., Araujo, L., Amann, J., Horn, L., … Carbone, D. P. (2014). Oncogenic and sorafenib-sensitive ARAF mutations in lung adenocarcinoma.Journal of Clinical Investigation,124(4), 1582–1586. https://doi.org/10.1172/JCI72763 |
| S214F | A-RAF | Yes | Between C1_1 and Tyrosine Protein Kinase | Gain-of-functi on | Imielinski, M., Greulich, H., Kaplan, B., Araujo, L., Amann, J., Horn, L., … Carbone, D. P. (2014). Oncogenic and sorafenib-sensitive ARAF mutations in lung adenocarcinoma.Journal of Clinical Investigation,124(4), 1582–1586. https://doi.org/10.1172/JCI72763 |
| S214T | A-RAF | Likely | Between C1_1 and Tyrosine Protein Kinase | Gain-of-functi on | Imielinski, M., Greulich, H., Kaplan, B., Araujo, L., Amann, J., Horn, L., … Carbone, D. P. (2014). Oncogenic and sorafenib-sensitive ARAF mutations in lung adenocarcinoma.Journal of Clinical Investigation,124(4), 1582–1586. https://doi.org/10.1172/JCI72763 |
| P216A | A-RAF | Likely | Between C1_1 and Tyrosine Protein Kinase | Likely Gain-of-functi on | Imielinski, M., Greulich, H., Kaplan, B., Araujo, L., Amann, J., Horn, L., … Carbone, D. P. (2014). Oncogenic and sorafenib-sensitive ARAF mutations in lung adenocarcinoma.Journal of Clinical Investigation,124(4), 1582–1586. https://doi.org/10.1172/JCI72763 |
| N217I | A-RAF | Yes | Between C1_1 and Tyrosine Protein Kinase | Gain-of-functi on | Sia, D., Losic, B., Moeini, A., Cabellos, L., Hao, K., Revill, K., … Llovet, J. M. (2015). Massive parallel sequencing uncovers actionable FGFR2–PPHLN1 fusion and ARAF mutations in intrahepatic cholangiocarcinoma.Nature Communications, 6(1), 6087. https://doi.org/10.1038/ncomms7087 |
| G322S | A-RAF | Likely | Tyrosine Protein Kinase | Gain-of-functi on | Sia, D., Losic, B., Moeini, A., Cabellos, L., Hao, K., Revill, K., … Llovet, J. M. (2015). Massive parallel sequencing uncovers actionable FGFR2–PPHLN1 fusion |

| | | | | | |
|---|---|---|---|---|---|
| | | | | | and ARAF mutations in intrahepatic cholangiocarcinoma.Nature Communications, 6(1), 6087. https://doi.org/10.1038/ncomms7087 |
| Q347_A348del | A-RAF | Yes | Tyrosine Protein Kinase | Gain-of-function | Nelson, D. S., Quispel, W., Badalian-Very, G., van Halteren, A. G. S., van den Bos, C., Bovee, J. V. M. G., … Rollins, B. J. (2014). Somatic activating ARAF mutations in Langerhans cell histiocytosis.Blood, 123(20), 3152–3155. https://doi.org/10.1182/blood-2013-06-511139 |
| F351L | A-RAF | Yes | Tyrosine Protein Kinase | Gain-of-function | Nelson, D. S., Quispel, W., Badalian-Very, G., van Halteren, A. G. S., van den Bos, C., Bovee, J. V. M. G., … Rollins, B. J. (2014). Somatic activating ARAF mutations in Langerhans cell histiocytosis.Blood, 123(20), 3152–3155. https://doi.org/10.1182/blood-2013-06-511139 |
| S151A | B-RAF | Likely | Before Ras-binding domain domain | Likely Gain-of-function | Jacobsen, E., Shanmugam, V., & Jagannathan, J. (2017). Rosai–Dorfman Disease with Activating KRAS Mutation — Response to Cobimetinib.New England Journal of Medicine, 377(24), 2398–2399. https://doi.org/10.1056/NEJMc1713676 |
| Q201H | B-RAF | Inconclusive | Ras-binding domain | Inconclusive | Estep, A. L., Palmer, C., McCormick, F., & Rauen, K. A. (2007). Mutation Analysis of BRAF, MEK1 and MEK2 in 15 Ovarian Cancer Cell Lines: Implications for Therapy.PLoS ONE, 2(12), e1279. https://doi.org/10.1371/journal.pone.0001279 |
| p61BRAF-V600E | B-RAF | Yes | Ras-binding domain | Gain-of-function | Poulikakos, P. I., Zhang, C., Bollag, G., Shokat, K. M., & Rosen, N. (2010). RAF inhibitors transactivate RAF dimers and ERK signalling in cells with wild-type BRAF. Nature, 464(7287), 427–430. https://doi.org/10.1038/nature08902 |
| T241P | B-RAF | Likely | C1_1 | Likely Gain-of-function | Sarkozy, A., Carta, C., Moretti, S., Zampino, G., Digilio, M. C., Pantaleoni, F., … Tartaglia, M. (2009). Germline BRAF mutations in Noonan, LEOPARD, and cardiofaciocutaneous syndromes: Molecular diversity and associated phenotypic spectrum.Human Mutation, 30(4), 695–702. https://doi.org/10.1002/humu.20955 |
| A246P | B-RAF | Likely | C1_1 | Gain-of-function | Niihori, T., Aoki, Y., Narumi, Y., Neri, G., Cavé, H., Verloes, A., … Matsubara, Y. (2006). Germline KRAS and BRAF mutations in cardio-facio-cutaneous syndrome.Nature Genetics, 38(3), 294–296. https://doi.org/10.1038/ng1749 |

| | | | | | |
|---|---|---|---|---|---|
| F247L | B-RAF | Likely | C1_1 | Likely Gain-of-function | Lu, H., Villafane, N., Dogruluk, T., Grzeskowiak, C. L., Kong, K., Tsang, Y. H., … Scott, K. L. (2017). Engineering and Functional Characterization of Fusion Genes Identifies Novel Oncogenic Drivers of Cancer. Cancer Research, 77(13), 3502–3512. https://doi.org/10.1158/0008-5472.CAN-16-2745 |
| Q257R | B-RAF | Likely | C1_1 | Gain-of-function | Anastasaki, C., Estep, A. L., Marais, R., Rauen, K. A., & Patton, E. E. (2009). Kinase-activating and kinase-impaired cardio-facio-cutaneous syndrome alleles have activity during zebrafish development and are sensitive to small molecule inhibitors.Human Molecular Genetics, 18(14), 2543–2554. https://doi.org/10.1093/hmg/ddp186 |
| E275K | B-RAF | Likely | C1_1 | Likely Gain-of-function | Sarkozy, A., Carta, C., Moretti, S., Zampino, G., Digilio, M. C., Pantaleoni, F., … Tartaglia, M. (2009). Germline BRAF mutations in Noonan, LEOPARD, and cardiofaciocutaneous syndromes: Molecular diversity and associated phenotypic spectrum.Human Mutation, 30(4), 695–702. https://doi.org/10.1002/humu.20955 |
| D287H | B-RAF | Yes | Between C1_1 and Tyrosine Protein Kinase | Gain-of-function | Yao, Z., Torres, N. M., Tao, A., Gao, Y., Luo, L., Li, Q., … Rosen, N. (2015). BRAF Mutants Evade ERK-Dependent Feedback by Different Mechanisms that Determine Their Sensitivity to Pharmacologic Inhibition.Cancer Cell, 28(3), 370–383. https://doi.org/10.1016/j.ccell.2015.08.001 |
| I326V | B-RAF | Likely Neutral | Between C1_1 and Tyrosine Protein Kinase | Likely Neutral | Razzaque, M. A., Nishizawa, T., Komoike, Y., Yagi, H., Furutani, M., Amo, R., … Matsuoka, R. (2007). Germline gain-of-function mutations in RAF1 cause Noonan syndrome.Nature Genetics, 39(8), 1013–1017. https://doi.org/10.1038/ng2078 |
| P367R | B-RAF | Likely | Between C1_1 and Tyrosine Protein Kinase | Likely Gain-of-function | Berger, A. H., Brooks, A. N., Wu, X., Shrestha, Y., Chouinard, C., Piccioni, F., … Boehm, J. S. (2016). High-throughput Phenotyping of Lung Cancer Somatic Mutations.Cancer Cell, 30(2), 214–228. https://doi.org/10.1016/j.ccell.2016.06.022 |
| W450L | B-RAF | Likely Neutral | Between C1_1 and Tyrosine Protein Kinase | Likely Neutral | Berger, A. H., Brooks, A. N., Wu, X., Shrestha, Y., Chouinard, C., Piccioni, F., … Boehm, J. S. (2016). High-throughput Phenotyping of Lung Cancer Somatic Mutations.Cancer Cell, 30(2), 214–228. https://doi.org/10.1016/j.ccell.2016.06.022 |
| V459L | B-RAF | Yes | Tyrosine Protein Kinase | Gain-of-function | Yao, Z., Yaeger, R., Rodrik-Outmezguine, V. S., Tao, A., Torres, N. M., Chang, M. T., … Rosen, N. (2017). Tumours with class 3 BRAF mutants are sensitive to the |

| | | | | | |
|---|---|---|---|---|---|
| | | | | | inhibition of activated RAS.Nature, 548(7666), 234–238. https://doi.org/10.1038/nature23291 |
| R462E | B-RAF | Likely | Tyrosine Protein Kinase | Likely Gain-of-functi on | Haling, J. R., Sudhamsu, J., Yen, I., Sideris, S., Sandoval, W., Phung, W., … Malek, S. (2014). Structure of the BRAF-MEK Complex Reveals a Kinase Activity Independent Role for BRAF in MAPK Signaling.Cancer Cell, 26(3), 402–413. https://doi.org/10.1016/j.ccr.2014.07.007 |
| R462I | B-RAF | Likely Neutral | Tyrosine Protein Kinase | Likely Neutral | Wan, P. T., Garnett, M. J., Roe, S. M., Lee, S., Niculescu-Duvaz, D., Good, V. M., ... & Marais, R. (2004). Mechanism of activation of the RAF-ERK signaling pathway by oncogenic mutations of B-RAF.Cell,116(6), 855-867. |
| I463S | B-RAF | Likely Neutral | Tyrosine Protein Kinase | Likely Neutral | Ikenoue, T., Hikiba, Y., Kanai, F., Aragaki, J., Tanaka, Y., Imamura, J., … Omata, M. (2004). Different effects of point mutations within the B-Raf glycine-rich loop in colorectal tumors on mitogen-activated protein/extracellular signal-regulated kinase kinase/extracellular signal-regulated kinase and nuclear factor kappaB pathway and cellular tra. Cancer Research, 64(10), 3428–3435. https://doi.org/10.1158/0008-5472.CAN-03-3591 |
| G464E | B-RAF | Yes | Tyrosine Protein Kinase | Gain-of-functi on | Ikenoue, T., Hikiba, Y., Kanai, F., Aragaki, J., Tanaka, Y., Imamura, J., … Omata, M. (2004). Different effects of point mutations within the B-Raf glycine-rich loop in colorectal tumors on mitogen-activated protein/extracellular signal-regulated kinase kinase/extracellular signal-regulated kinase and nuclear factor kappaB pathway and cellular tra. Cancer Research, 64(10), 3428–3435. https://doi.org/10.1158/0008-5472.CAN-03-3591 |
| G464R | B-RAF | Likely | Tyrosine Protein Kinase | Gain-of-functi on | Houben, R., Becker, J. C., Kappel, A., Terheyden, P., Bröcker, E.-B., Goetz, R., & Rapp, U. R. (2004). Constitutive activation of the Ras-Raf signaling pathway in metastatic melanoma is associated with poor prognosis. Journal of Carcinogenesis, 3, 6. https://doi.org/10.1186/1477-3163-3-6 |
| G464V | B-RAF | Yes | Tyrosine Protein Kinase | Gain-of-functi on | Ikenoue, T., Hikiba, Y., Kanai, F., Aragaki, J., Tanaka, Y., Imamura, J., … Omata, M. (2004). Different effects of point mutations within the B-Raf glycine-rich loop in colorectal tumors on mitogen-activated protein/extracellular signal-regulated kinase kinase/extracellular signal-regulated kinase and nuclear factor kappaB pathway and cellular tra. Cancer Research, 64(10), 3428–3435. https://doi.org/10.1158/0008-5472.CAN-03-3591 |

| | | | | | |
|---|---|---|---|---|---|
| G466A | B-RAF | Yes | Tyrosine Protein Kinase | Gain-of-functi on | Yao, Z., Yaeger, R., Rodrik-Outmezguine, V. S., Tao, A., Torres, N. M., Chang, M. T., … Rosen, N. (2017). Tumours with class 3 BRAF mutants are sensitive to the inhibition of activated RAS. Nature, 548(7666), 234–238. https://doi.org/10.1038/nature23291 |
| G466E | B-RAF | Yes | Tyrosine Protein Kinase | Gain-of-functi on | Richtig, G., Aigelsreiter, A., Kashofer, K., Talakic, E., Kupsa, R., Schaider, H., & Richtig, E. (n.d.). Two Case Reports of Rare BRAF Mutations in Exon 11 and Exon 15 with Discussion of Potential Treatment Options. Case Reports in Oncology, 9(3), 543–546. https://doi.org/10.1159/000449125 |
| G466R | B-RAF | Likely | Tyrosine Protein Kinase | Likely Gain-of-functi on | Garnett, M. J., Rana, S., Paterson, H., Barford, D., & Marais, R. (2005). Wild-type and mutant B-RAF activate C-RAF through distinct mechanisms involving heterodimerization. Molecular Cell, 20(6), 963–969. https://doi.org/10.1016/j.molcel.2005.10.022 |
| G466V | B-RAF | Yes | Tyrosine Protein Kinase | Gain-of-functi on | Sen, B., Peng, S., Tang, X., Erickson, H. S., Galindo, H., Mazumdar, T., … Johnson, F. M. (2012). Kinase-impaired BRAF mutations in lung cancer confer sensitivity to dasatinib. Science Translational Medicine, 4(136), 136ra70. https://doi.org/10.1126/scitranslmed.3003513 |
| S467A | B-RAF | Likely | Tyrosine Protein Kinase | Gain-of-functi on | Anastasaki, C., Estep, A. L., Marais, R., Rauen, K. A., & Patton, E. E. (2009). Kinase-activating and kinase-impaired cardio-facio-cutaneous syndrome alleles have activity during zebrafish development and are sensitive to small molecule inhibitors. Human Molecular Genetics, 18(14), 2543–2554. https://doi.org/10.1093/hmg/ddp186 |
| S467L | B-RAF | Yes | Tyrosine Protein Kinase | Gain-of-functi on | Zheng, G., Tseng, L.-H., Chen, G., Haley, L., Illei, P., Gocke, C. D., … Lin, M.-T. (2015). Clinical detection and categorization of uncommon and concomitant mutations involving BRAF. BMC Cancer, 15, 779. https://doi.org/10.1186/s12885-015-1811-y |
| F468C | B-RAF | Yes | Tyrosine Protein Kinase | Gain-of-functi on | Ikenoue, T., Hikiba, Y., Kanai, F., Aragaki, J., Tanaka, Y., Imamura, J., … Omata, M. (2004). Different effects of point mutations within the B-Raf glycine-rich loop in colorectal tumors on mitogen-activated protein/extracellular signal-regulated kinase kinase/extracellular signal-regulated kinase and nuclear factor kappaB pathway and cellular tra. Cancer Research, 64(10), 3428–3435. https://doi.org/10.1158/0008-5472.CAN-03-3591 |

| | | | | | |
|---|---|---|---|---|---|
| G469A | B-RAF | Yes | Tyrosine Protein Kinase | Gain-of-functi on | Poulikakos, P. I., Zhang, C., Bollag, G., Shokat, K. M., & Rosen, N. (2010). RAF inhibitors transactivate RAF dimers and ERK signalling in cells with wild-type BRAF. Nature, 464(7287), 427–430. https://doi.org/10.1038/nature08902 |
| G469del | B-RAF | Likely | Tyrosine Protein Kinase | Likely Gain-of-functi on | Cardarella, S., Ogino, A., Nishino, M., Butaney, M., Shen, J., Lydon, C., … Jänne, P. A. (2013). Clinical, pathologic, and biologic features associated with BRAF mutations in non-small cell lung cancer. Clinical Cancer Research : An Official Journal of the American Association for Cancer Research, 19(16), 4532–4540. https://doi.org/10.1158/1078-0432.CCR-13-0657 |
| G469E | B-RAF | Yes | Tyrosine Protein Kinase | Gain-of-functi on | Ikenoue, T., Hikiba, Y., Kanai, F., Aragaki, J., Tanaka, Y., Imamura, J., … Omata, M. (2004). Different effects of point mutations within the B-Raf glycine-rich loop in colorectal tumors on mitogen-activated protein/extracellular signal-regulated kinase kinase/extracellular signal-regulated kinase and nuclear factor kappaB pathway and cellular tra. Cancer Research, 64(10), 3428–3435. https://doi.org/10.1158/0008-5472.CAN-03-3591 |
| G469L | B-RAF | Likely | Tyrosine Protein Kinase | Likely Gain-of-functi on | Damm, F., Mylonas, E., Cosson, A., Yoshida, K., Della Valle, V., Mouly, E., … Bernard, O. A. (2014). Acquired initiating mutations in early hematopoietic cells of CLL patients. Cancer Discovery, 4(9), 1088–1101. https://doi.org/10.1158/2159-8290.CD-14-0104 |
| G469R | B-RAF | Yes | Tyrosine Protein Kinase | Gain-of-functi on | Hussain, M. R. M., Baig, M., Mohamoud, H. S. A., Ulhaq, Z., Hoessli, D. C., Khogeer, G. S., … Al-Aama, J. Y. (2015). BRAF gene: From human cancers to developmental syndromes. Saudi Journal of Biological Sciences, 22(4), 359–373. https://doi.org/10.1016/j.sjbs.2014.10.002 |
| G469V | B-RAF | Yes | Tyrosine Protein Kinase | Gain-of-functi on | Casadei Gardini, A., Chiadini, E., Faloppi, L., Marisi, G., Delmonte, A., Scartozzi, M., … Ulivi, P. (2016). Efficacy of sorafenib in BRAF-mutated non-small-cell lung cancer (NSCLC) and no response in synchronous BRAF wild type-hepatocellular carcinoma: a case report. BMC Cancer, 16, 429. https://doi.org/10.1186/s12885-016-2463-2 |
| V471F | B-RAF | Yes | Tyrosine Protein Kinase | Gain-of-functi on | Hu, J., Ahuja, L. G., Meharena, H. S., Kannan, N., Kornev, A. P., Taylor, S. S., & Shaw, A. S. (2015). Kinase regulation by hydrophobic spine assembly in cancer. Molecular and Cellular Biology, 35(1), 264–276. https://doi.org/10.1128/MCB.00943-14 |

| | | | | | |
|---|---|---|---|---|---|
| Y472C | B-RAF | Likely | Tyrosine Protein Kinase | Gain-of-function | Sen, B., Peng, S., Tang, X., Erickson, H. S., Galindo, H., Mazumdar, T., … Johnson, F. M. (2012). Kinase-impaired BRAF mutations in lung cancer confer sensitivity to dasatinib. Science Translational Medicine, 4(136), 136ra70. https://doi.org/10.1126/scitranslmed.3003513 |
| G478C | B-RAF | Likely Neutral | Tyrosine Protein Kinase | Loss-of-function | Rebocho, A. P., & Marais, R. (2013). ARAF acts as a scaffold to stabilize BRAF:CRAF heterodimers. Oncogene, 32(26), 3207–3212. https://doi.org/10.1038/onc.2012.330 |
| K483E | B-RAF | Likely | Tyrosine Protein Kinase | Likely Gain-of-function | Dela Cruz, F. S., Diolaiti, D., Turk, A. T., Rainey, A. R., Ambesi-Impiombato, A., Andrews, S. J., … Kung, A. L. (2016). A case study of an integrative genomic and experimental therapeutic approach for rare tumors: identification of vulnerabilities in a pediatric poorly differentiated carcinoma. Genome Medicine, 8(1), 116. https://doi.org/10.1186/s13073-016-0366-0 |
| K483M | B-RAF | Likely | Tyrosine Protein Kinase | Likely Gain-of-function | Anastasaki, C., Estep, A. L., Marais, R., Rauen, K. A., & Patton, E. E. (2009). Kinase-activating and kinase-impaired cardio-facio-cutaneous syndrome alleles have activity during zebrafish development and are sensitive to small molecule inhibitors. Human Molecular Genetics, 18(14), 2543–2554. https://doi.org/10.1093/hmg/ddp186 |
| L485_P490del | B-RAF | Yes | Tyrosine Protein Kinase | Gain-of-function | Chen, S., Zhang, Y., Van Horn, R., Yin, T., Buchanan, S., Yadav, V., … Peng, S. (2016). Oncogenic BRAF Deletions That Function as Homodimers and Are Sensitive to Inhibition by RAF Dimer Inhibitor LY3009120. Cancer Discovery, 6(3), 300–315. https://doi.org/10.1158/2159-8290 |
| L485_P490delinsF | B-RAF | Likely | Tyrosine Protein Kinase | Likely Gain-of-function | Chen, S., Zhang, Y., Van Horn, R., Yin, T., Buchanan, S., Yadav, V., … Peng, S. (2016). Oncogenic BRAF Deletions That Function as Homodimers and Are Sensitive to Inhibition by RAF Dimer Inhibitor LY3009120. Cancer Discovery, 6(3), 300–315. https://doi.org/10.1158/2159-8290 |
| L485_P490delinsY | B-RAF | Likely | Tyrosine Protein Kinase | Likely Gain-of-function | Chen, S., Zhang, Y., Van Horn, R., Yin, T., Buchanan, S., Yadav, V., … Peng, S. (2016). Oncogenic BRAF Deletions That Function as Homodimers and Are Sensitive to Inhibition by RAF Dimer Inhibitor LY3009120. Cancer Discovery, 6(3), 300–315. https://doi.org/10.1158/2159-8290 |

| | | | | | |
|---|---|---|---|---|---|
| L485_Q494del | B-RAF | Likely | Tyrosine Protein Kinase | Likely Gain-of-function | Chen, S., Zhang, Y., Van Horn, R., Yin, T., Buchanan, S., Yadav, V., … Peng, S. (2016). Oncogenic BRAF Deletions That Function as Homodimers and Are Sensitive to Inhibition by RAF Dimer Inhibitor LY3009120. Cancer Discovery, 6(3), 300–315. https://doi.org/10.1158/2159-8290 |
| L485F | B-RAF | Likely | Tyrosine Protein Kinase | Gain-of-function | Niihori, T., Aoki, Y., Narumi, Y., Neri, G., Cavé, H., Verloes, A., … Matsubara, Y. (2006). Germline KRAS and BRAF mutations in cardio-facio-cutaneous syndrome. Nature Genetics, 38(3), 294–296. https://doi.org/10.1038/ng1749 |
| N486_P490del | B-RAF | Likely | Tyrosine Protein Kinase | Likely Gain-of-function | Chen, S., Zhang, Y., Van Horn, R., Yin, T., Buchanan, S., Yadav, V., … Peng, S. (2016). Oncogenic BRAF Deletions That Function as Homodimers and Are Sensitive to Inhibition by RAF Dimer Inhibitor LY3009120. Cancer Discovery, 6(3), 300–315. https://doi.org/10.1158/2159-8290 |
| N486_T491delinsK | B-RAF | Likely | Tyrosine Protein Kinase | Likely Gain-of-function | Diamond, E. L., Durham, B. H., Ulaner, G. A., Drill, E., Buthorn, J., Ki, M., … Hyman, D. M. (2019). Efficacy of MEK inhibition in patients with histiocytic neoplasms. Nature, 567(7749), 521–524. https://doi.org/10.1038/s41586-019-1012-y |
| V487_P492delinsA | B-RAF | Likely | Tyrosine Protein Kinase | Likely Gain-of-function | Chen, S., Zhang, Y., Van Horn, R., Yin, T., Buchanan, S., Yadav, V., … Peng, S. (2016). Oncogenic BRAF Deletions That Function as Homodimers and Are Sensitive to Inhibition by RAF Dimer Inhibitor LY3009120. Cancer Discovery, 6(3), 300–315. https://doi.org/10.1158/2159-8290 |
| T488_P492del | B-RAF | Likely | Tyrosine Protein Kinase | Likely Gain-of-function | Chen, S., Zhang, Y., Van Horn, R., Yin, T., Buchanan, S., Yadav, V., … Peng, S. (2016). Oncogenic BRAF Deletions That Function as Homodimers and Are Sensitive to Inhibition by RAF Dimer Inhibitor LY3009120. Cancer Discovery, 6(3), 300–315. https://doi.org/10.1158/2159-8290 |
| P490_Q494del | B-RAF | Likely | Tyrosine Protein Kinase | Likely Gain-of-function | Chen, S., Zhang, Y., Van Horn, R., Yin, T., Buchanan, S., Yadav, V., … Peng, S. (2016). Oncogenic BRAF Deletions That Function as Homodimers and Are Sensitive to Inhibition by RAF Dimer Inhibitor LY3009120. Cancer Discovery, 6(3), 300–315. https://doi.org/10.1158/2159-8290 |
| K499E | B-RAF | Likely | Tyrosine Protein Kinase | Gain-of-function | Niihori, T., Aoki, Y., Narumi, Y., Neri, G., Cavé, H., Verloes, A., … Matsubara, Y. (2006). Germline KRAS and BRAF mutations in cardio-facio-cutaneous syndrome. Nature Genetics, 38(3), 294–296. https://doi.org/10.1038/ng1749 |

| | | | | | |
|---|---|---|---|---|---|
| E501G | B-RAF | Likely Neutral | Tyrosine Protein Kinase | Likely Loss-of-function | Niihori, T., Aoki, Y., Narumi, Y., Neri, G., Cavé, H., Verloes, A., … Matsubara, Y. (2006). Germline KRAS and BRAF mutations in cardio-facio-cutaneous syndrome. Nature Genetics, 38(3), 294–296. https://doi.org/10.1038/ng1749 |
| E501K | B-RAF | Inconclusive | Tyrosine Protein Kinase | Inconclusive | Razzaque, M. A., Nishizawa, T., Komoike, Y., Yagi, H., Furutani, M., Amo, R., … Matsuoka, R. (2007). Germline gain-of-function mutations in RAF1 cause Noonan syndrome. Nature Genetics, 39(8), 1013–1017. https://doi.org/10.1038/ng2078 |
| L505H | B-RAF | Yes | Tyrosine Protein Kinase | Gain-of-function | Hoogstraat, M., Gadellaa-van Hooijdonk, C. G., Ubink, I., Besselink, N. J. M., Pieterse, M., Veldhuis, W., … Lolkema, M. P. (2015). Detailed imaging and genetic analysis reveal a secondary BRAF(L505H) resistance mutation and extensive intrapatient heterogeneity in metastatic BRAF mutant melanoma patients treated with vemurafenib. Pigment Cell & Melanoma Research, 28(3), 318–323. https://doi.org/10.1111/pcmr.12347 |
| R506_K507insVLR | B-RAF | Likely | Tyrosine Protein Kinase | Gain-of-function | Jones, D. T. W., Hutter, B., Jäger, N., Korshunov, A., Kool, M., Warnatz, H.-J., … International Cancer Genome Consortium PedBrain Tumor Project. (2013). Recurrent somatic alterations of FGFR1 and NTRK2 in pilocytic astrocytoma. Nature Genetics, 45(8), 927–932. https://doi.org/10.1038/ng.2682 |
| R509H | B-RAF | Likely Neutral | Tyrosine Protein Kinase | Loss-of-function | Poulikakos, P. I., Zhang, C., Bollag, G., Shokat, K. M., & Rosen, N. (2010). RAF inhibitors transactivate RAF dimers and ERK signalling in cells with wild-type BRAF. Nature, 464(7287), 427–430. https://doi.org/10.1038/nature08902 |
| L514V | B-RAF | Likely | Tyrosine Protein Kinase | Likely Gain-of-function | Wang, J., Yao, Z., Jonsson, P., Allen, A. N., Qin, A. C. R., Uddin, S., … Pratilas, C. A. (2018). A Secondary Mutation in BRAF Confers Resistance to RAF Inhibition in a BRAFV600E-Mutant Brain Tumor. Cancer Discovery, 8(9), 1130–1141. https://doi.org/10.1158/2159-8290.CD-17-1263 |
| T529I | B-RAF | Likely Neutral | Tyrosine Protein Kinase | Likely Neutral | Daub, H., Specht, K., & Ullrich, A. (2004). Strategies to overcome resistance to targeted protein kinase inhibitors. Nature Reviews. Drug Discovery, 3(12), 1001–1010. https://doi.org/10.1038/nrd1579 |
| T529M | B-RAF | Likely Neutral | Tyrosine Protein Kinase | Likely Neutral | Daub, H., Specht, K., & Ullrich, A. (2004). Strategies to overcome resistance to targeted protein kinase inhibitors. Nature Reviews. Drug Discovery, 3(12), 1001–1010. https://doi.org/10.1038/nrd1579 |

| | | | | | |
|---|---|---|---|---|---|
| T529N | B-RAF | Likely Neutral | Tyrosine Protein Kinase | Likely Neutral | Daub, H., Specht, K., & Ullrich, A. (2004). Strategies to overcome resistance to targeted protein kinase inhibitors. Nature Reviews. Drug Discovery, 3(12), 1001–1010. https://doi.org/10.1038/nrd1579 |
| W531C | B-RAF | Likely Neutral | Tyrosine Protein Kinase | Likely Neutral | Sarkozy, A., Carta, C., Moretti, S., Zampino, G., Digilio, M. C., Pantaleoni, F., … Tartaglia, M. (2009). Germline BRAF mutations in Noonan, LEOPARD, and cardiofaciocutaneous syndromes: Molecular diversity and associated phenotypic spectrum. Human Mutation, 30(4), 695–702. https://doi.org/10.1002/humu.20955 |
| H574Q | B-RAF | Likely | Tyrosine Protein Kinase | Likely Gain-of-functi on | Berger, A. H., Brooks, A. N., Wu, X., Shrestha, Y., Chouinard, C., Piccioni, F., … Boehm, J. S. (2016). High-throughput Phenotyping of Lung Cancer Somatic Mutations. Cancer Cell, 30(2), 214–228. https://doi.org/10.1016/j.ccell.2016.06.022 |
| N581D | B-RAF | Likely | Tyrosine Protein Kinase | Likely Gain-of-functi on | Niihori, T., Aoki, Y., Narumi, Y., Neri, G., Cavé, H., Verloes, A., … Matsubara, Y. (2006). Germline KRAS and BRAF mutations in cardio-facio-cutaneous syndrome. Nature Genetics, 38(3), 294–296. https://doi.org/10.1038/ng1749 |
| N581I | B-RAF | Yes | Tyrosine Protein Kinase | Gain-of-functi on | Wong, C. W., Fan, Y. S., Chan, T. L., Chan, A. S. W., Ho, L. C., Ma, T. K. F., … Cancer Genome Project. (2005). BRAF and NRAS mutations are uncommon in melanomas arising in diverse internal organs. Journal of Clinical Pathology, 58(6), 640–644. https://doi.org/10.1136/jcp.2004.022509 |
| N581S | B-RAF | Yes | Tyrosine Protein Kinase | Gain-of-functi on | Yao, Z., Yaeger, R., Rodrik-Outmezguine, V. S., Tao, A., Torres, N. M., Chang, M. T., … Rosen, N. (2017). Tumours with class 3 BRAF mutants are sensitive to the inhibition of activated RAS. Nature, 548(7666), 234–238. https://doi.org/10.1038/nature23291 |
| N581Y | B-RAF | Likely | Tyrosine Protein Kinase | Likely Gain-of-functi on | Seth, R., Crook, S., Ibrahem, S., Fadhil, W., Jackson, D., & Ilyas, M. (2009). Concomitant mutations and splice variants in KRAS and BRAF demonstrate complex perturbation of the Ras/Raf signalling pathway in advanced colorectal cancer. Gut, 58(9), 1234–1241. https://doi.org/10.1136/gut.2008.159137 |
| L584F | B-RAF | Inconclusive | Tyrosine Protein Kinase | Inconclusive | Stones, C. J., Kim, J. E., Joseph, W. R., Leung, E., Marshall, E. S., Finlay, G. J., … Baguley, B. C. (2013). Comparison of responses of human melanoma cell lines to MEK and BRAF inhibitors. Frontiers in Genetics, 4, 66. https://doi.org/10.3389/fgene.2013.00066 |

| | | | | | |
|---|---|---|---|---|---|
| E586K | B-RAF | Likely | Tyrosine Protein Kinase | Gain-of-functi on | Wan, P. T. C., Garnett, M. J., Roe, S. M., Lee, S., Niculescu-Duvaz, D., Good, V. M., … Cancer Genome Project. (2004). Mechanism of activation of the RAF-ERK signaling pathway by oncogenic mutations of B-RAF. Cell, 116(6), 855–867. https://doi.org/10.1016/s0092-8674(04)00215-6 |
| D594A | B-RAF | Yes | Tyrosine Protein Kinase | Gain-of-functi on | Yao, Z., Yaeger, R., Rodrik-Outmezguine, V. S., Tao, A., Torres, N. M., Chang, M. T., … Rosen, N. (2017). Tumours with class 3 BRAF mutants are sensitive to the inhibition of activated RAS. Nature, 548(7666), 234–238. https://doi.org/10.1038/nature23291 |
| D594E | B-RAF | Likely | Tyrosine Protein Kinase | Gain-of-functi on | Kamata, T., Hussain, J., Giblett, S., Hayward, R., Marais, R., & Pritchard, C. (2010). BRAF inactivation drives aneuploidy by deregulating CRAF. Cancer Research, 70(21), 8475–8486. https://doi.org/10.1158/0008-5472.CAN-10-0603 |
| D594G | B-RAF | Yes | Tyrosine Protein Kinase | Gain-of-functi on | Zheng, G., Tseng, L.-H., Chen, G., Haley, L., Illei, P., Gocke, C. D., … Lin, M.-T. (2015). Clinical detection and categorization of uncommon and concomitant mutations involving BRAF. BMC Cancer, 15, 779. https://doi.org/10.1186/s12885-015-1811-y |
| D594H | B-RAF | Yes | Tyrosine Protein Kinase | Gain-of-functi on | Yao, Z., Yaeger, R., Rodrik-Outmezguine, V. S., Tao, A., Torres, N. M., Chang, M. T., … Rosen, N. (2017). Tumours with class 3 BRAF mutants are sensitive to the inhibition of activated RAS. Nature, 548(7666), 234–238. https://doi.org/10.1038/nature23291 |
| D594N | B-RAF | Yes | Tyrosine Protein Kinase | Gain-of-functi on | Cardarella, S., Ogino, A., Nishino, M., Butaney, M., Shen, J., Lydon, C., … Jänne, P. A. (2013). Clinical, pathologic, and biologic features associated with BRAF mutations in non-small cell lung cancer. Clinical Cancer Research : An Official Journal of the American Association for Cancer Research, 19(16), 4532–4540. https://doi.org/10.1158/1078-0432.CCR-13-0657 |
| D594V | B-RAF | Likely | Tyrosine Protein Kinase | Gain-of-functi on | Ikenoue, T., Hikiba, Y., Kanai, F., Tanaka, Y., Imamura, J., Imamura, T., … Omata, M. (2003). Functional analysis of mutations within the kinase activation segment of B-Raf in human colorectal tumors. Cancer Research, 63(23), 8132–8137. Retrieved from http://www.ncbi.nlm.nih.gov/pubmed/14678966 |

| | | | | | |
|---|---|---|---|---|---|
| D594Y | B-RAF | Likely | Tyrosine Protein Kinase | Likely Gain-of-functi on | Cardarella, S., Ogino, A., Nishino, M., Butaney, M., Shen, J., Lydon, C., … Jänne, P. A. (2013). Clinical, pathologic, and biologic features associated with BRAF mutations in non-small cell lung cancer. Clinical Cancer Research : An Official Journal of the American Association for Cancer Research, 19(16), 4532–4540. https://doi.org/10.1158/1078-0432.CCR-13-0657 |
| F595L | B-RAF | Yes | Tyrosine Protein Kinase | Gain-of-functi on | Ikenoue, T., Hikiba, Y., Kanai, F., Aragaki, J., Tanaka, Y., Imamura, J., … Omata, M. (2004). Different effects of point mutations within the B-Raf glycine-rich loop in colorectal tumors on mitogen-activated protein/extracellular signal-regulated kinase kinase/extracellular signal-regulated kinase and nuclear factor kappaB pathway and cellular tra. Cancer Research, 64(10), 3428–3435. https://doi.org/10.1158/0008-5472.CAN-03-3591 |
| G596C | B-RAF | Likely | Tyrosine Protein Kinase | Gain-of-functi on | Noeparast, A., Teugels, E., Giron, P., Verschelden, G., De Brakeleer, S., Decoster, L., & De Grève, J. (2017). Non-V600 BRAF mutations recurrently found in lung cancer predict sensitivity to the combination of Trametinib and Dabrafenib. Oncotarget, 8(36). https://doi.org/10.18632/oncotarget.11635 |
| G596D | B-RAF | Yes | Tyrosine Protein Kinase | Gain-of-functi on | Kang, S., Kim, H.-S., Seo, S. S., Park, S.-Y., Sidransky, D., & Dong, S. M. (2007). Inverse correlation between RASSF1A hypermethylation, KRAS and BRAF mutations in cervical adenocarcinoma. Gynecologic Oncology, 105(3), 662–666. https://doi.org/10.1016/j.ygyno.2007.01.045 |
| G596R | B-RAF | Yes | Tyrosine Protein Kinase | Gain-of-functi on | Ikenoue, T., Hikiba, Y., Kanai, F., Tanaka, Y., Imamura, J., Imamura, T., … Omata, M. (2003). Functional analysis of mutations within the kinase activation segment of B-Raf in human colorectal tumors. Cancer Research, 63(23), 8132–8137. Retrieved from http://www.ncbi.nlm.nih.gov/pubmed/14678966 |
| G596V | B-RAF | Likely | Tyrosine Protein Kinase | Likely Gain-of-functi on | Anastasaki, C., Estep, A. L., Marais, R., Rauen, K. A., & Patton, E. E. (2009). Kinase-activating and kinase-impaired cardio-facio-cutaneous syndrome alleles have activity during zebrafish development and are sensitive to small molecule inhibitors. Human Molecular Genetics, 18(14), 2543–2554. https://doi.org/10.1093/hmg/ddp186 |
| L597Q | B-RAF | Yes | Tyrosine Protein Kinase | Gain-of-functi on | Poulikakos, P. I., Zhang, C., Bollag, G., Shokat, K. M., & Rosen, N. (2010). RAF inhibitors transactivate RAF dimers and ERK signalling in cells with wild-type BRAF. Nature, 464(7287), 427–430. https://doi.org/10.1038/nature08902 |

| | | | | | |
|---|---|---|---|---|---|
| L597R | B-RAF | Likely | Tyrosine Protein Kinase | Likely Gain-of-function | Dahlman, K. B., Xia, J., Hutchinson, K., Ng, C., Hucks, D., Jia, P., … Pao, W. (2012). BRAF(L597) mutations in melanoma are associated with sensitivity to MEK inhibitors. Cancer Discovery, 2(9), 791–797. https://doi.org/10.1158/2159-8290.CD-12-0097 |
| L597S | B-RAF | Likely | Tyrosine Protein Kinase | Likely Gain-of-function | Dahlman, K. B., Xia, J., Hutchinson, K., Ng, C., Hucks, D., Jia, P., … Pao, W. (2012). BRAF(L597) mutations in melanoma are associated with sensitivity to MEK inhibitors. Cancer Discovery, 2(9), 791–797. https://doi.org/10.1158/2159-8290.CD-12-0097 |
| L597V | B-RAF | Yes | Tyrosine Protein Kinase | Gain-of-function | Poulikakos, P. I., Zhang, C., Bollag, G., Shokat, K. M., & Rosen, N. (2010). RAF inhibitors transactivate RAF dimers and ERK signalling in cells with wild-type BRAF. Nature, 464(7287), 427–430. https://doi.org/10.1038/nature08902 |
| A598T | B-RAF | Likely Neutral | Tyrosine Protein Kinase | Loss-of-function | Chang, M. T., Asthana, S., Gao, S. P., Lee, B. H., Chapman, J. S., Kandoth, C., … Taylor, B. S. (2016). Identifying recurrent mutations in cancer reveals widespread lineage diversity and mutational specificity. Nature Biotechnology, 34(2), 155–163. https://doi.org/10.1038/nbt.3391 |
| A598V | B-RAF | Likely | Tyrosine Protein Kinase | Gain-of-function | Santarpia, L., Sherman, S. I., Marabotti, A., Clayman, G. L., & El-Naggar, A. K. (2009). Detection and molecular characterization of a novel BRAF activated domain mutation in follicular variant of papillary thyroid carcinoma. Human Pathology, 40(6), 827–833. https://doi.org/10.1016/j.humpath.2008.11.003 |
| T599_V600insEAT | B-RAF | Likely | Tyrosine Protein Kinase | Likely Gain-of-function | Moretti, S., Macchiarulo, A., De Falco, V., Avenia, N., Barbi, F., Carta, C., … Puxeddu, E. (2006). Biochemical and molecular characterization of the novel BRAF(V599Ins) mutation detected in a classic papillary thyroid carcinoma. Oncogene, 25(30), 4235–4240. https://doi.org/10.1038/sj.onc.1209448 |
| T599_V600insETT | B-RAF | Likely | Tyrosine Protein Kinase | Likely Gain-of-function | Moretti, S., Macchiarulo, A., De Falco, V., Avenia, N., Barbi, F., Carta, C., … Puxeddu, E. (2006). Biochemical and molecular characterization of the novel BRAF(V599Ins) mutation detected in a classic papillary thyroid carcinoma. Oncogene, 25(30), 4235–4240. https://doi.org/10.1038/sj.onc.1209448 |
| T599_V600insV | B-RAF | Yes | Tyrosine Protein Kinase | Gain-of-function | Moretti, S., Macchiarulo, A., De Falco, V., Avenia, N., Barbi, F., Carta, C., … Puxeddu, E. (2006). Biochemical and molecular characterization of the novel BRAF(V599Ins) mutation detected in a classic papillary thyroid carcinoma. Oncogene, 25(30), 4235–4240. https://doi.org/10.1038/sj.onc.1209448 |

| | | | | | |
|---|---|---|---|---|---|
| T599dup | B-RAF | Yes | Tyrosine Protein Kinase | Gain-of-functi on | Cardarella, S., Ogino, A., Nishino, M., Butaney, M., Shen, J., Lydon, C., … Jänne, P. A. (2013). Clinical, pathologic, and biologic features associated with BRAF mutations in non-small cell lung cancer. Clinical Cancer Research : An Official Journal of the American Association for Cancer Research, 19(16), 4532–4540. https://doi.org/10.1158/1078-0432.CCR-13-0657 |
| T599I | B-RAF | Likely | Tyrosine Protein Kinase | Gain-of-functi on | Wan, P. T. C., Garnett, M. J., Roe, S. M., Lee, S., Niculescu-Duvaz, D., Good, V. M., … Cancer Genome Project. (2004). Mechanism of activation of the RAF-ERK signaling pathway by oncogenic mutations of B-RAF. Cell, 116(6), 855–867. https://doi.org/10.1016/s0092-8674(04)00215-6 |
| T599insTT | B-RAF | Yes | Tyrosine Protein Kinase | Gain-of-functi on | Eisenhardt, A. E., Olbrich, H., Röring, M., Janzarik, W., Anh, T. N. Van, Cin, H., … Brummer, T. (2011). Functional characterization of a BRAF insertion mutant associated with pilocytic astrocytoma. International Journal of Cancer, 129(9), 2297–2303. https://doi.org/10.1002/ijc.25893 |
| T599R | B-RAF | Yes | Tyrosine Protein Kinase | Gain-of-functi on | Cho, U., Oh, W. J., Bae, J. S., Lee, S., Lee, Y. S., Park, G. S., … Jung, C. K. (2014). Clinicopathological Features of Rare BRAF Mutations in Korean Thyroid Cancer Patients. Journal of Korean Medical Science, 29(8), 1054. https://doi.org/10.3346/jkms.2014.29.8.1054 |
| V600_K601de linsE | B-RAF | Yes | Tyrosine Protein Kinase | Gain-of-functi on | El Karak, F., Assi, T., Kourie, H. R., El Rassy, E., Chebib, R., Ghor, M., & Tabchi, S. (2015). BRAFV600 mutant non-small-cell lung cancer resistant to Vemurafenib. International Journal of Clinical and Experimental Pathology, 8(3), 3294–3298. Retrieved from http://www.ncbi.nlm.nih.gov/pubmed/26045855 |
| V600D | B-RAF | Yes | Tyrosine Protein Kinase | Gain-of-functi on | Cancer Genome Atlas Research Network. (2014). Integrated genomic characterization of papillary thyroid carcinoma. Cell, 159(3), 676–690. https://doi.org/10.1016/j.cell.2014.09.050 |
| V600D_K601i nsFGLAT | B-RAF | Yes | Tyrosine Protein Kinase | Gain-of-functi on | Hou, P., Liu, D., & Xing, M. (2007). Functional characterization of the T1799-1801del and A1799-1816ins BRAF mutations in papillary thyroid cancer. Cell Cycle (Georgetown, Tex.), 6(3), 377–379. https://doi.org/10.4161/cc.6.3.3818 |
| V600delinsY M | B-RAF | Yes | Tyrosine Protein Kinase | Gain-of-functi on | Matsuse, M., Mitsutake, N., Tanimura, S., Ogi, T., Nishihara, E., Hirokawa, M., … Yamashita, S. (2013). Functional characterization of the novel BRAF complex mutation, BRAF(V600delinsYM) , identified in papillary thyroid carcinoma. |

| | | | | | |
|---|---|---|---|---|---|
| | | | | | International Journal of Cancer, 132(3), 738–743. https://doi.org/10.1002/ijc.27709 |
| V600E | B-RAF | Yes | Tyrosine Protein Kinase | Gain-of-functi on | Cancer Genome Atlas Research Network. (2014). Integrated genomic characterization of papillary thyroid carcinoma. Cell, 159(3), 676–690. https://doi.org/10.1016/j.cell.2014.09.050 |
| V600G | B-RAF | Likely | Tyrosine Protein Kinase | Gain-of-functi on | Champion, K. J., Bunag, C., Estep, A. L., Jones, J. R., Bolt, C. H., Rogers, R. C., … Everman, D. B. (2011). Germline mutation in BRAF codon 600 is compatible with human development: de novo p.V600G mutation identified in a patient with CFC syndrome. Clinical Genetics, 79(5), 468–474. https://doi.org/10.1111/j.1399-0004.2010.01495.x |
| V600K | B-RAF | Yes | Tyrosine Protein Kinase | Gain-of-functi on | Cancer Genome Atlas Research Network. (2014). Integrated genomic characterization of papillary thyroid carcinoma. Cell, 159(3), 676–690. https://doi.org/10.1016/j.cell.2014.09.050 |
| V600M | B-RAF | Likely | Tyrosine Protein Kinase | Gain-of-functi on | Heinzerling, L., Kühnapfel, S., Meckbach, D., Baiter, M., Kaempgen, E., Keikavoussi, P., … Schneider-Stock, R. (2013). Rare BRAF mutations in melanoma patients: implications for molecular testing in clinical practice. British Journal of Cancer, 108(10), 2164–2171. https://doi.org/10.1038/bjc.2013.143 |
| V600R | B-RAF | Yes | Tyrosine Protein Kinase | Gain-of-functi on | Cancer Genome Atlas Research Network. (2014). Integrated genomic characterization of papillary thyroid carcinoma. Cell, 159(3), 676–690. https://doi.org/10.1016/j.cell.2014.09.050 |
| K601E | B-RAF | Likely | Tyrosine Protein Kinase | Gain-of-functi on | Poulikakos, P. I., Zhang, C., Bollag, G., Shokat, K. M., & Rosen, N. (2010). RAF inhibitors transactivate RAF dimers and ERK signalling in cells with wild-type BRAF. Nature, 464(7287), 427–430. https://doi.org/10.1038/nature08902 |
| K601N | B-RAF | Yes | Tyrosine Protein Kinase | Gain-of-functi on | Poulikakos, P. I., Zhang, C., Bollag, G., Shokat, K. M., & Rosen, N. (2010). RAF inhibitors transactivate RAF dimers and ERK signalling in cells with wild-type BRAF. Nature, 464(7287), 427–430. https://doi.org/10.1038/nature08902 |
| K601Q | B-RAF | Likely | Tyrosine Protein Kinase | Gain-of-functi on | Sarkozy, A., Carta, C., Moretti, S., Zampino, G., Digilio, M. C., Pantaleoni, F., … Tartaglia, M. (2009). Germline BRAF mutations in Noonan, LEOPARD, and cardiofaciocutaneous syndromes: Molecular diversity and associated phenotypic spectrum. Human Mutation, 30(4), 695–702. https://doi.org/10.1002/humu.20955 |

| | | | | | |
|---|---|---|---|---|---|
| K601T | B-RAF | Yes | Tyrosine Protein Kinase | Gain-of-functi on | Lionetti, M., Barbieri, M., Todoerti, K., Agnelli, L., Marzorati, S., Fabris, S., … Neri, A. (2015). Molecular spectrum of BRAF, NRAS and KRAS gene mutations in plasma cell dyscrasias: implication for MEK-ERK pathway activation. Oncotarget, 6(27), 24205–24217. https://doi.org/10.18632/oncotarget.4434 |
| R671Q | B-RAF | Likely | Tyrosine Protein Kinase | Likely Gain-of-functi on | Wan, L., Chen, M., Cao, J., Dai, X., Yin, Q., Zhang, J., … Wei, W. (2017). The APC/C E3 Ligase Complex Activator FZR1 Restricts BRAF Oncogenic Function. Cancer Discovery, 7(4), 424–441. https://doi.org/10.1158/2159-8290.CD-16-0647 |
| A728V | B-RAF | Likely | After Tyrosine Protein Kinase | Gain-of-functi on | Wan, P. T. C., Garnett, M. J., Roe, S. M., Lee, S., Niculescu-Duvaz, D., Good, V. M., … Cancer Genome Project. (2004). Mechanism of activation of the RAF-ERK signaling pathway by oncogenic mutations of B-RAF. Cell, 116(6), 855–867. https://doi.org/10.1016/s0092-8674(04)00215-6 |
| E104K | C-RAF | Likely Neutral | Ras-binding domain | Likely Neutral | Antony, R., Emery, C. M., Sawyer, A. M., & Garraway, L. A. (2013). C-RAF mutations confer resistance to RAF inhibitors. Cancer Research, 73(15), 4840–4851. https://doi.org/10.1158/0008-5472.CAN-12-4089 |
| K106N | C-RAF | Likely | Ras-binding domain | Likely Gain-of-functi on | Diamond, E. L., Durham, B. H., Ulaner, G. A., Drill, E., Buthorn, J., Ki, M., … Hyman, D. M. (2019). Efficacy of MEK inhibition in patients with histiocytic neoplasms. Nature, 567(7749), 521–524. https://doi.org/10.1038/s41586-019-1012-y |
| S257L | C-RAF | Yes | Between C1_1 and Tyrosine Protein Kinase | Gain-of-functi on | Light, Y., Paterson, H., & Marais, R. (2002). 14-3-3 antagonizes Ras-mediated Raf-1 recruitment to the plasma membrane to maintain signaling fidelity. Molecular and Cellular Biology, 22(14), 4984–4996. https://doi.org/10.1128/mcb.22.14.4984-4996.2002 |
| S257P | C-RAF | Likely Neutral | Between C1_1 and Tyrosine Protein Kinase | Likely Neutral | Antony, R., Emery, C. M., Sawyer, A. M., & Garraway, L. A. (2013). C-RAF mutations confer resistance to RAF inhibitors. Cancer Research, 73(15), 4840–4851. https://doi.org/10.1158/0008-5472.CAN-12-4089 |
| S257W | C-RAF | Likely | Between C1_1 and Tyrosine Protein Kinase | Gain-of-functi on | Light, Y., Paterson, H., & Marais, R. (2002). 14-3-3 antagonizes Ras-mediated Raf-1 recruitment to the plasma membrane to maintain signaling fidelity. Molecular and Cellular Biology, 22(14), 4984–4996. https://doi.org/10.1128/mcb.22.14.4984-4996.2002 |

| | | | | | |
|---|---|---|---|---|---|
| S259A | C-RAF | Yes | Between C1_1 and Tyrosine Protein Kinase | Gain-of-functi on | Imielinski, M., Greulich, H., Kaplan, B., Araujo, L., Amann, J., Horn, L., … Carbone, D. P. (2014). Oncogenic and sorafenib-sensitive ARAF mutations in lung adenocarcinoma. Journal of Clinical Investigation, 124(4), 1582–1586. https://doi.org/10.1172/JCI72763 |
| S259F | C-RAF | Yes | Between C1_1 and Tyrosine Protein Kinase | Gain-of-functi on | Pandit, B., Sarkozy, A., Pennacchio, L. A., Carta, C., Oishi, K., Martinelli, S., … Gelb, B. D. (2007). Gain-of-function RAF1 mutations cause Noonan and LEOPARD syndromes with hypertrophic cardiomyopathy. Nature Genetics, 39(8), 1007–1012. https://doi.org/10.1038/ng2073 |
| S259P | C-RAF | Likely | Between C1_1 and Tyrosine Protein Kinase | Gain-of-functi on | Molzan, M., Schumacher, B., Ottmann, C., Baljuls, A., Polzien, L., Weyand, M., … Ottmann, C. (2010). Impaired binding of 14-3-3 to C-RAF in Noonan syndrome suggests new approaches in diseases with increased Ras signaling. Molecular and Cellular Biology, 30(19), 4698–4711. https://doi.org/10.1128/MCB.01636-09 |
| P261L | C-RAF | Likely | Between C1_1 and Tyrosine Protein Kinase | Likely Gain-of-functi on | Molzan, M., Schumacher, B., Ottmann, C., Baljuls, A., Polzien, L., Weyand, M., … Ottmann, C. (2010). Impaired binding of 14-3-3 to C-RAF in Noonan syndrome suggests new approaches in diseases with increased Ras signaling. Molecular and Cellular Biology, 30(19), 4698–4711. https://doi.org/10.1128/MCB.01636-09 |
| P261S | C-RAF | Likely | Between C1_1 and Tyrosine Protein Kinase | Likely Gain-of-functi on | Pandit, B., Sarkozy, A., Pennacchio, L. A., Carta, C., Oishi, K., Martinelli, S., … Gelb, B. D. (2007). Gain-of-function RAF1 mutations cause Noonan and LEOPARD syndromes with hypertrophic cardiomyopathy. Nature Genetics, 39(8), 1007–1012. https://doi.org/10.1038/ng2073 |
| P261T | C-RAF | Likely | Between C1_1 and Tyrosine Protein Kinase | Likely Gain-of-functi on | Antony, R., Emery, C. M., Sawyer, A. M., & Garraway, L. A. (2013). C-RAF mutations confer resistance to RAF inhibitors. Cancer Research, 73(15), 4840–4851. https://doi.org/10.1158/0008-5472.CAN-12-4089 |
| R318W | C-RAF | Inconclusive | Between C1_1 and Tyrosine Protein Kinase | Inconclusive | Eisfeld, A.-K., Kohlschmidt, J., Mrózek, K., Volinia, S., Blachly, J. S., Nicolet, D., … Bloomfield, C. D. (2017). Mutational Landscape and Gene Expression Patterns in Adult Acute Myeloid Leukemias with Monosomy 7 as a Sole Abnormality. Cancer Research, 77(1), 207–218. https://doi.org/10.1158/0008-5472.CAN-16-1386 |

| | | | | | |
|---|---|---|---|---|---|
| G356E | C-RAF | Likely | Tyrosine Protein Kinase | Likely Gain-of-functi on | Antony, R., Emery, C. M., Sawyer, A. M., & Garraway, L. A. (2013). C-RAF mutations confer resistance to RAF inhibitors. Cancer Research, 73(15), 4840–4851. https://doi.org/10.1158/0008-5472.CAN-12-4089 |
| G361A | C-RAF | Likely | Tyrosine Protein Kinase | Likely Gain-of-functi on | Antony, R., Emery, C. M., Sawyer, A. M., & Garraway, L. A. (2013). C-RAF mutations confer resistance to RAF inhibitors. Cancer Research, 73(15), 4840–4851. https://doi.org/10.1158/0008-5472.CAN-12-4089 |
| R391W | C-RAF | Likely | Tyrosine Protein Kinase | Likely Gain-of-functi on | Atefi, M., Titz, B., Tsoi, J., Avramis, E., Le, A., Ng, C., … Graeber, T. G. (2016). CRAF R391W is a melanoma driver oncogene. Scientific Reports, 6, 27454. https://doi.org/10.1038/srep27454 |
| S427G | C-RAF | Yes | Tyrosine Protein Kinase | Gain-of-functi on | Zebisch, A., Staber, P. B., Delavar, A., Bodner, C., Hiden, K., Fischereder, K., … Sill, H. (2006). Two transforming C-RAF germ-line mutations identified in patients with therapy-related acute myeloid leukemia. Cancer Research, 66(7), 3401–3408. https://doi.org/10.1158/0008-5472.CAN-05-0115 |
| S427T | C-RAF | Likely Neutral | Tyrosine Protein Kinase | Likely Neutral | Antony, R., Emery, C. M., Sawyer, A. M., & Garraway, L. A. (2013). C-RAF mutations confer resistance to RAF inhibitors. Cancer Research, 73(15), 4840–4851. https://doi.org/10.1158/0008-5472.CAN-12-4089 |
| D447N | C-RAF | Likely | Tyrosine Protein Kinase | Likely Gain-of-functi on | Antony, R., Emery, C. M., Sawyer, A. M., & Garraway, L. A. (2013). C-RAF mutations confer resistance to RAF inhibitors. Cancer Research, 73(15), 4840–4851. https://doi.org/10.1158/0008-5472.CAN-12-4089 |
| I448V | C-RAF | Yes | Tyrosine Protein Kinase | Gain-of-functi on | Zebisch, A., Haller, M., Hiden, K., Goebel, T., Hoefler, G., Troppmair, J., & Sill, H. (2009). Loss of RAF kinase inhibitor protein is a somatic event in the pathogenesis of therapy-related acute myeloid leukemias with C-RAF germline mutations. Leukemia, 23(6), 1049–1053. https://doi.org/10.1038/leu.2009.68 |
| M469I | C-RAF | Likely | Tyrosine Protein Kinase | Likely Gain-of-functi on | Antony, R., Emery, C. M., Sawyer, A. M., & Garraway, L. A. (2013). C-RAF mutations confer resistance to RAF inhibitors. Cancer Research, 73(15), 4840–4851. https://doi.org/10.1158/0008-5472.CAN-12-4089 |
| E478K | C-RAF | Likely | Tyrosine Protein Kinase | Likely Gain-of-functi on | Antony, R., Emery, C. M., Sawyer, A. M., & Garraway, L. A. (2013). C-RAF mutations confer resistance to RAF inhibitors. Cancer Research, 73(15), 4840–4851. https://doi.org/10.1158/0008-5472.CAN-12-4089 |

| | | | | | |
|---|---|---|---|---|---|
| D486N | C-RAF | Likely | Tyrosine Protein Kinase | Likely Gain-of-functi on | Wu, X., Yin, J., Simpson, J., Kim, K.-H., Gu, S., Hong, J. H., … Araki, T. (2012). Increased BRAF heterodimerization is the common pathogenic mechanism for noonan syndrome-associated RAF1 mutants. Molecular and Cellular Biology, 32(19), 3872–3890. https://doi.org/10.1128/MCB.00751-12 |
| R554K | C-RAF | Likely Neutral | Tyrosine Protein Kinase | Likely Neutral | Antony, R., Emery, C. M., Sawyer, A. M., & Garraway, L. A. (2013). C-RAF mutations confer resistance to RAF inhibitors. Cancer Research, 73(15), 4840–4851. https://doi.org/10.1158/0008-5472.CAN-12-4089 |
| L613V | C-RAF | Likely | After Tyrosine Protein Kinase | Likely Gain-of-functi on | Wu, X., Yin, J., Simpson, J., Kim, K.-H., Gu, S., Hong, J. H., … Araki, T. (2012). Increased BRAF heterodimerization is the common pathogenic mechanism for noonan syndrome-associated RAF1 mutants. Molecular and Cellular Biology, 32(19), 3872–3890. https://doi.org/10.1128/MCB.00751-12 |

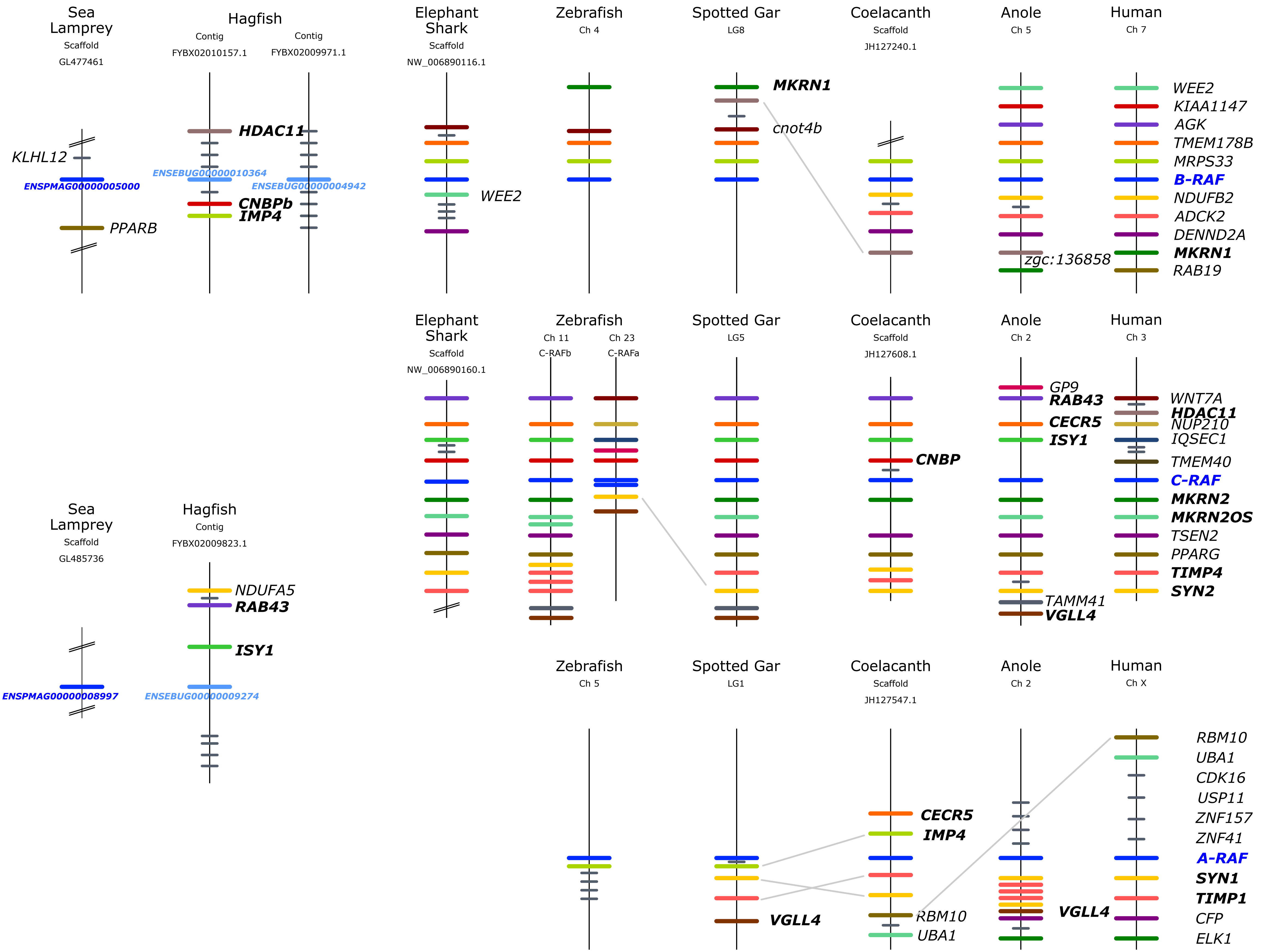
Sea Lamprey
Scaffold
GL477461
KLHL12
ENSPMAG00000005000
PPARB
Hagfish
Contig
FYBX02010157.1
Contig
FYBX02009971.1
HDAC11
ENSEBUG00000010364
ENSEBUG00000004942
CNBPb
IMP4
Elephant Shark
Scaffold
NW_006890116.1
WEE2
Zebrafish
Ch 4
Spotted Gar
LG8
MKRN1
cnot4b
Coelacanth
Scaffold
JH127240.1
Anole
Ch 5
zgc:136858
Human
Ch 7
WEE2
KIAA1147
AGK
TMEM178B
MRPS33
B-RAF
NDUFB2
ADCK2
DENND2A
MKRN1
RAB19
Elephant Shark
Scaffold
NW_006890160.1
Zebrafish
Ch 11
C-RAFb
Ch 23
C-RAFa
Spotted Gar
LG5
Coelacanth
Scaffold
JH127608.1
CNBP
Anole
Ch 2
GP9
RAB43
CECR5
ISY1
TAMM41
VGLL4
Human
Ch 3
WNT7A
HDAC11
NUP210
IQSEC1
TMEM40
C-RAF
MKRN2
MKRN2OS
TSEN2
PPARG
TIMP4
SYN2
Sea Lamprey
Scaffold
GL485736
ENSPMAG00000008997
Hagfish
Contig
FYBX02009823.1
NDUFA5
RAB43
ISY1
ENSEBUG00000009274
Zebrafish
Ch 5
Spotted Gar
LG1
VGLL4
Coelacanth
Scaffold
JH127547.1
CECR5
IMP4
RBM10
UBA1
Anole
Ch 2
VGLL4
Human
Ch X
RBM10
UBA1
CDK16
USP11
ZNF157
ZNF41
A-RAF
SYN1
TIMP1
CFP
ELK1
